# DreamAI: algorithm for the imputation of proteomics data

**DOI:** 10.1101/2020.07.21.214205

**Authors:** Weiping Ma, Sunkyu Kim, Shrabanti Chowdhury, Zhi Li, Mi Yang, Seungyeul Yoo, Francesca Petralia, Jeremy Jacobsen, Jingyi Jessica Li, Xinzhou Ge, Kexin Li, Thomas Yu, Anna P. Calinawan, Nathan Edwards, Samuel H. Payne, Paul C. Boutros, Henry Rodriguez, Gustavo Stolovitzky, Jun Zhu, Jaewoo Kang, David Fenyo, Julio Saez-Rodriguez, Pei Wang

**Author notes:** Corresponding author: Pei Wang. Co-first authors (Weiping Ma, Sunkyu Kim).

## Abstract

Deep proteomics profiling using labeled LC-MS/MS experiments has been proven to be powerful to study complex diseases. However, due to the dynamic nature of the discovery mass spectrometry, the generated data contain a substantial fraction of missing values. This poses great challenges for data analyses, as many tools, especially those for high dimensional data, cannot deal with missing values directly. To address this problem, the NCI-CPTAC Proteogenomics DREAM Challenge was carried out to develop effective imputation algorithms for labeled LC-MS/MS proteomics data through crowd learning. The final resulting algorithm, DreamAI, is based on an ensemble of six different imputation methods. The imputation accuracy of DreamAI, as measured by Pearson correlation, is about 15%-50% greater than existing tools among less abundant proteins, which are more vulnerable to be missed in proteomics data sets. This new tool notably enhances data analysis capabilities in proteomics research.

## Introduction

Proteins are responsible for nearly every task of cellular life and are important molecules for disease diagnosis, prevention and treatment. The technique of Liquid Chromatography Tandem Mass Spectrometry (LC-MS/MS) using isobaric labeling methods, including isobaric tags for absolute and relative quantification (iTRAQ) and tandem mass tags (TMT), allows detection and quantification of thousands of proteins and tens of thousands of their post-translational modifications (PTM) in a given biological sample [1,2]. Isobaric labeling not only greatly enhances the precision of quantification, but also improves the throughput [3,4], as multiple samples can be combined into one multiplex and profiled simultaneously. These technological developments greatly accelerate the application of proteomics in various diseases studies [1,2,5,6,7,8]. However, missingness in mass spectrometry (MS) based proteomics can limit the usability of this data.

Only a subset of peptides and PTMs in a biological sample can be detected and quantified in each LC-MS/MS experiment, due to a number of factors, including: the proteome complexity of many biological samples; the stochastic sampling procedure; and the limited duty cycle of mass spectrometry based discovery proteomics. The members of this detectable subset vary from experiment to experiment. Thus, when proteomic profiles from a collection of LC-MS/MS experiments are analyzed together, a substantial number of missing values are present [9]. In addition, the multiplex structure of isobaric labeling experiments affects the missingness rate, since the detection of a peptide is performed for all samples in MS1 within the multiplex. Consequently, a peptide is either observed or missing for all samples analyzed together in one multiplex. This type of experimentally-induced multiplex-level missing mechanism constitutes the majority of missing events when using isobaric labeling. For example, in proteomics data sets generated in CPTAC/TCGA Ovarian Cancer Study with iTRAQ platform[2], among all detected proteins and phospho-sites, 31.1% proteins and 98.3% phospho-sites had missing values in at least one sample (**Fig. 1a-b, Fig. S1a-b**). And more than 95% or 99% of total missing events in the global or phospho-proteomics data sets are multiplex-level missing (**Fig. 1c**). This multiplex-level missingness is also prevalent in data from TMT platforms, as illustrated in **Fig. 1a-b** and **S1a-b** based on data examples from the CPTAC Prospective Ovarian Cancer Study [7].

**Figure 1.**
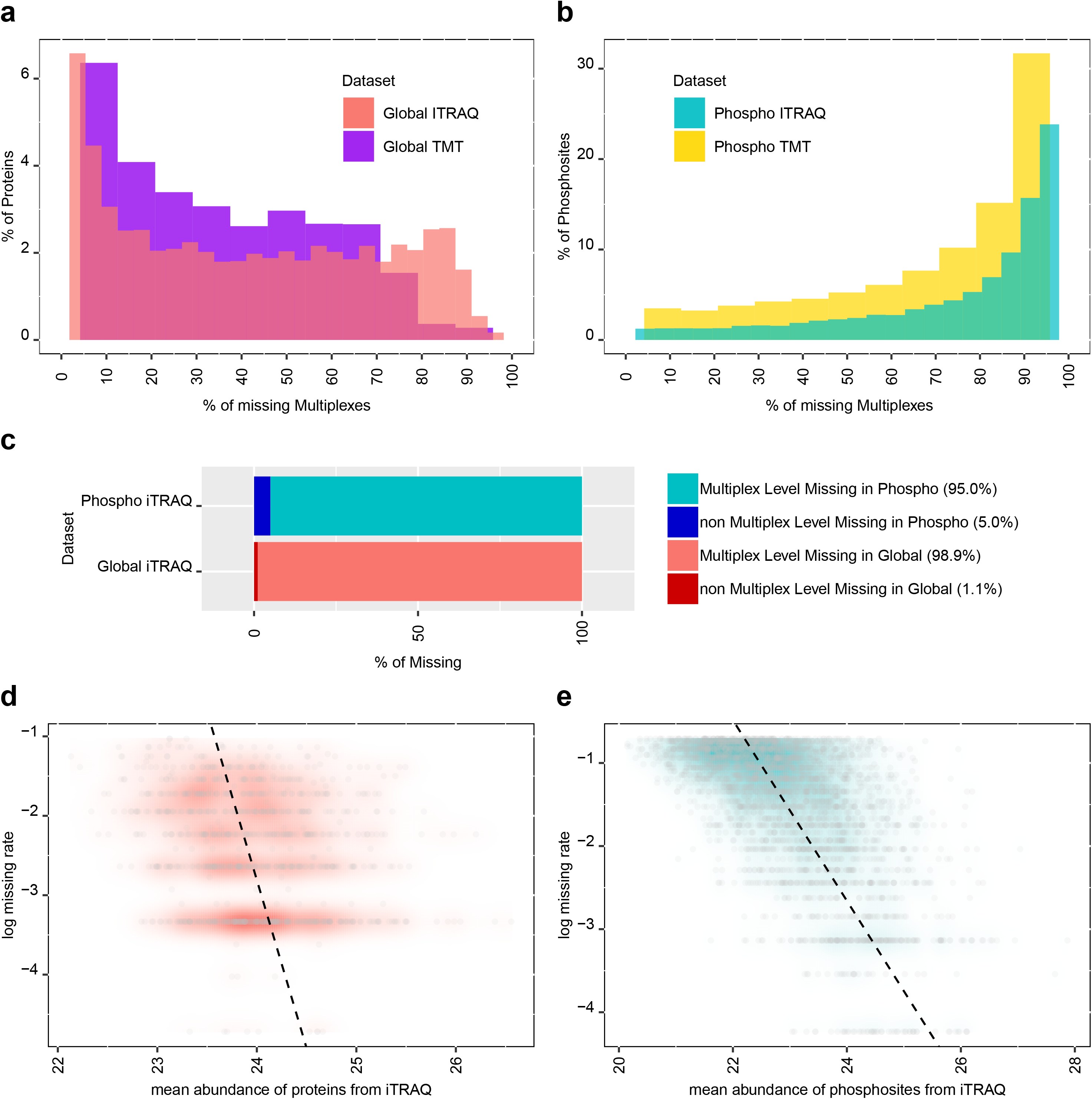
Missing rates and missing patterns in various proteomics data sets from CPTAC ovarian studies [2,7]. **(a)** Distribution of protein level missing rates in a 4plex-iTRAQ global-proteomics data set of 112 tumor samples [2] and a 10plex-TMT global proteomics data set of 120 tumor samples [7]. X-axis represents the percentage of iTRAQ/TMT multiplexes in which individual proteins were not identified/quantified **(b)** Distribution of phospho-site level missing rates in a 4plex-iTRAQ phospho-proteomics data set of 92 samples and a 10plex-TMT phospho-proteomics data set of 120 samples. X-axis represents the percentage of iTRAQ/TMT multiplexes in which individual phosphosites were not identified/quantified. **(c)** Percentage of multiplex-level and non-multiplex-level missing data in iTRAQ global- and phospho-proteomics data sets. **(d)** Scatter plot of protein-level missing rates v.s. mean protein abundances based on observed data in the iTRAQ global-proteomics data set. **(e)** Scatter plot of phospho-site level missing rates v.s. mean phospho-site abundances based on observed data in the iTRAQ phospho-proteomics data set.

Missingness in mass spectrometry (MS) based proteomics data could be caused by many technical factors. As suggested in previous works [10,11,12], in labeled proteomics experiments, failures of the mass spectrometer to detect a peptide can be due to low peptide abundance, low peptide-dependent ionization efficiency, different distributions across multiple peptide charge states, and different modified forms of the peptide [13]. These factors all contribute to the abundance dependent missing tendency, wherein peptides/proteins with lower abundances tend to have higher probabilities of being missed, as illustrated in **Fig. 1d-e, and S1c-d** using MS1 (parent ion) intensity based qualification (please see **Supplement A3** for quantification details). Unsurprisingly, the degree of this dependence between missing probability and the abundance to be measured in labeled proteomics data varies across different experiments and studies, as many factors mentioned above work together to cause the missingness. The dependence between the propensity of missing on data points and the underlying values is referred to as MNAR --- *missing not at random* [14]. It has been well established in the statistical literature that, in the presence of MNAR, analysis based on the observed data only shall lead to biased estimates and incorrect inference [14].

The substantial missing rates combined with multiplex dependent MNAR bring great challenges to the downstream data analysis. The simply strategy of focusing only on proteins observed in all samples [1,2] makes the downstream data analysis convenient, but abandons a large amount of information from hundreds or thousands of proteins in each proteomics data set. Unfortunately, some very important proteins for understanding disease mechanisms could be abandoned in this process, as disease-relevant proteins are often low abundant or subtypes specific and therefore less likely to be measured in all samples.

Thus, there is a pressing need to have strategies other than simply ignoring proteins and PTMs with missing values in proteomics data analysis. Two commonly used methods for handling data with missing values are: 1. to substitute missing values with some constants (e.g., a small number or an estimated mean/median value) [15]; and 2. to perform analysis using observed data only [1,2]. The constant imputation, as well as its enhanced variation Perseus [16] which fills in missing values with random variables independently drawn from a pre-specified Gaussian distribution, do not work for labeled proteomics data, due to the experimentally induced multiplex-level missing patterns. On the other hand, for mass spectrometry data with MNAR, it is dangerous to perform analyses based on observed data points only, which could lead to biased estimates and incorrect inferences [10,14]. In addition, for multivariate and high-dimensional analysis, a subset of samples with completely observed features could be small or non-existent.

A more sensible solution is to perform stage-wise learning: firstly, use information from observed data points to “learn” the unobserved data points, i.e. impute the missing values; and then conduct statistical analysis based on the imputed matrices. Since proteins and PTMs that interact with each other usually have correlated abundances, the measured abundances in a given sample contain substantial information of other unobserved proteins and PTMs. Information from other samples with shared properties can also be useful in this learning step.

A few imputation strategies have been proposed to handle missing values in high dimension omics data sets in the past decades. Some of the strategies take advantage of local similarity within the data set. For example, the commonly used *KNN* imputation method predicts missing values based on information from K nearest neighbors (proteins or samples) [17,18](see **Supplement A8**). This strategy has been applied to a few proteogenomics studies [5]. In addition, *MissForest*, which builds Random Forest models to predict missing values of one feature based on observed values of all other features[19] (see **Supplement A8**), is another effective local similarity based imputation strategy and has been adopted in multiple genomics studies[20,21].

To better accommodate the MNAR in proteomics data, Chen et. al. (2014) [22] proposed an exponential function based probability model to characterize the abundance dependent missing pattern in unlabeled LC-MS/MS proteomics data, and a penalized EM algorithm (PEMM) to fit the model, which can be used for imputation[23]. To further accommodate the multiplex level missing structure in the labeled proteomics data, the authors further introduced mixEMM [10], which not only models the abundance dependent batch-level missing pattern but also utilizes mixed effects to better account for experimental variations across multiplexes. Based on mixEMM, in [6], the authors implemented an Abundance-Dependent Missing ImputatioN algorithm for labeled proteomics data [6], which essentially utilized a weighted average of neighboring data points to impute the missing values (see **Supplement A8**). This method is referred to as *ADMIN* in the rest of the paper.

Besides methods relying on local similarity in the data, there is a collection of imputation algorithms utilizing global structure of the data based on low rank matrix completion. Those methods that stemmed from the field of image de-noising[17,24,25,26], have flourished in a broad range of applications to solve various imputation problems, such as completion of single cell RNA-seq data[27] and GWAS data[28], as well as prediction of miRNA-Disease association[29]. Low rank matrix completion techniques have been recently applied to proteomics data imputation too. For example, *pcaMethods*, a principal component analysis (PCA) based method for matrix completion [30], has been applied to impute missing values in TMT proteomics data sets in a recent publication.[31]

Thorough efforts have been made to evaluate the performances of different imputation strategies on label-free proteomics data [12,32]. Consensus conclusions from these studies suggest that local similarity based methods and global structure based methods perform better than the constant imputation methods in the presence of MNAR [12,32]. In addition, one study [32] reported superior performance of methods based on global structure, such as low-rank matrix completion [18] and linear model based maximum likelihood estimate [33] [34] compared to those of local similarity based methods (KNN) for label free proteomics data. Moreover, as expected, it is more challenging to impute missing values for features with missing rate higher than 50% than those with lower missing rates [12].

Recently, pioneer efforts have also been made to study how various imputation tools work on labeled LC-MS/MS data sets. The investigation by Palstrøm et. al.[31] confirms the advantage of KNN and low rank matrix completion over constant imputation for labeled proteomics data. But the investigation is incomprehensive, due to the limited number of imputation methods considered and the inadequate numerical examples with rather simplified missing mechanism assumptions. In another recent review work [23], Bramer et. al. compared the performance of nine imputation algorithms on isobaric labeling-based proteomics data and found that PEMM and missForest have the best performances. However, most of the imputation methods considered in this investigation are not specifically developed for labeled LC-MS/MS data or even proteomics data. Therefore, it is of great interest to develop and systematically assess tools for imputing the missing values in proteomics data from labeled LC-MS/MS experiments.

Towards this goal, we carried out a NCI-CPTAC (Clinical Proteomic Tumor Analysis Consortium) DREAM Proteogenomics Imputation Challenge (https://sagebionetworks.org/research-projects/nci-cptac-dream-proteogenomics-challenge/). The challenge aimed to leverage techniques from multiple research fields such as statistical computation and machine learning, and to achieve a superior solution for the data imputation problem for labeled LC-MS/MS proteomics data sets through crowd learning.

The Challenge included a competition phase and a collaborative phase. In the competition phase, participants were invited to build imputation algorithms with training data sets and submit algorithms to test on additional data sets for evaluation and ranking. Training and testing data sets were simulated from the proteomics data sets in CPTAC breast cancer studies[1,8]. In the collaborative phase, together with the three winning teams from the competition phase, we further enhanced and integrated different imputation techniques. This effort led to the development of the final *Aggregation based Imputation algorithm* --- DreamAI, which is based on ensemble of six different imputation methods including: two low-rank matrix completion methods, two prediction based imputation methods, and two KNN type methods. The performance of DreamAI and other imputation tools were then systematically evaluated and compared using the CPTAC/TCGA Ovarian Cancer proteomics data sets, which contain profiles of duplicate tumor samples from the same patients [2]. The imputation accuracy of DreamAI, as measured by Pearson correlation coefficient (Cor), is about 15%-50% greater than the few leading popular tools, including ADMIN [6], KNN[17,18], missForest[19] and pcaMethods[30].

To illustrate the usage of imputation in proteomics data analysis, we performed proteogenomic integrative analysis using newly published data of deep TMT proteomics profiling of 103 clear cell renal cell carcinoma (CCRCC) samples and 80 adjacent normal tissue samples[35]. We observed improved RNA-protein concordances between transcriptomics and proteomics data after performing imputation on proteomics data. When evaluating the power to detect proteins having significantly different abundances between tumor and adjacent normal tissues, we further observed an advantage of DreamAI imputation as compared to mean- and KNN- imputation, as well as no imputation.

In summary, this work represents a landmark crowd-sourced community effort to address the problem of imputation for labeled LC-MS/MS proteomics data sets. The resulting source code, including an R package and Docker image of DreamAI are publicly available. This tool can benefit data analysis practice in a broad range of proteomics research.

## Experimental Design and Statistical Rationale

### Challenge overview

The NCI-CPTAC DREAM Proteogenomics Imputation Challenge was carried out to develop a benchmark imputation strategy for labeled LC-MS/MS proteomics data sets through crowd learning. The challenge consists of two phases: a challenging and a community phase. In the challenging phase, participants were invited to build their own imputation algorithms and winners were identified based on performances of submitted imputation algorithms on test data sets. In the community phase, the 3 top-performing teams worked jointly to develop a benchmark imputation strategy for labeled LC-MS/MS proteomics data.

### The challenging phase

Imputation is an unsupervised learning, which makes judging results particularly challenging. To objectively evaluate different imputation algorithms in the challenge phase, we implemented a simulation framework to generate decoy data sets with missing patterns mimicking that of the real data sets. We made use of the global proteomics profiles of labeled LC-MS/MS experiments from the CPTAC/TCGA breast cancer and ovarian cancer studies [1,7]. We started with the subsets of proteins with complete measurements and superimposed pseudo missing data points generated from probability models, mimicking the missing mechanisms in labeled proteomics experiments. We introduced both biological and instrumental missing events, with the probability of the latter depending on protein abundance measurements (see **Supplement A2** and **A3** for more details of data sets and data generation).

In total, 10 training data sets and 100 testing data sets were generated. The large number of test data sets was to allow a thorough evaluation of performances of submitted imputation algorithms (**Fig 2a, Supplement A4**). The training data sets were generated based on the global proteomics data from the CPTAC/TCGA Breast Cancer Study [1] and were shared with the challenge participants. The testing datasets were generated based on the global proteomics data from CPTAC Prospective Ovarian Cancer Study [7], which were not available to the public during the challenging period. Moreover, the simulated testing data sets were not shared with the participants. Each participating team needed to firstly develop an imputation algorithm based on training data sets, and then submit their final algorithm to Synapse to perform imputation on testing data sets.

**Figure 2.**
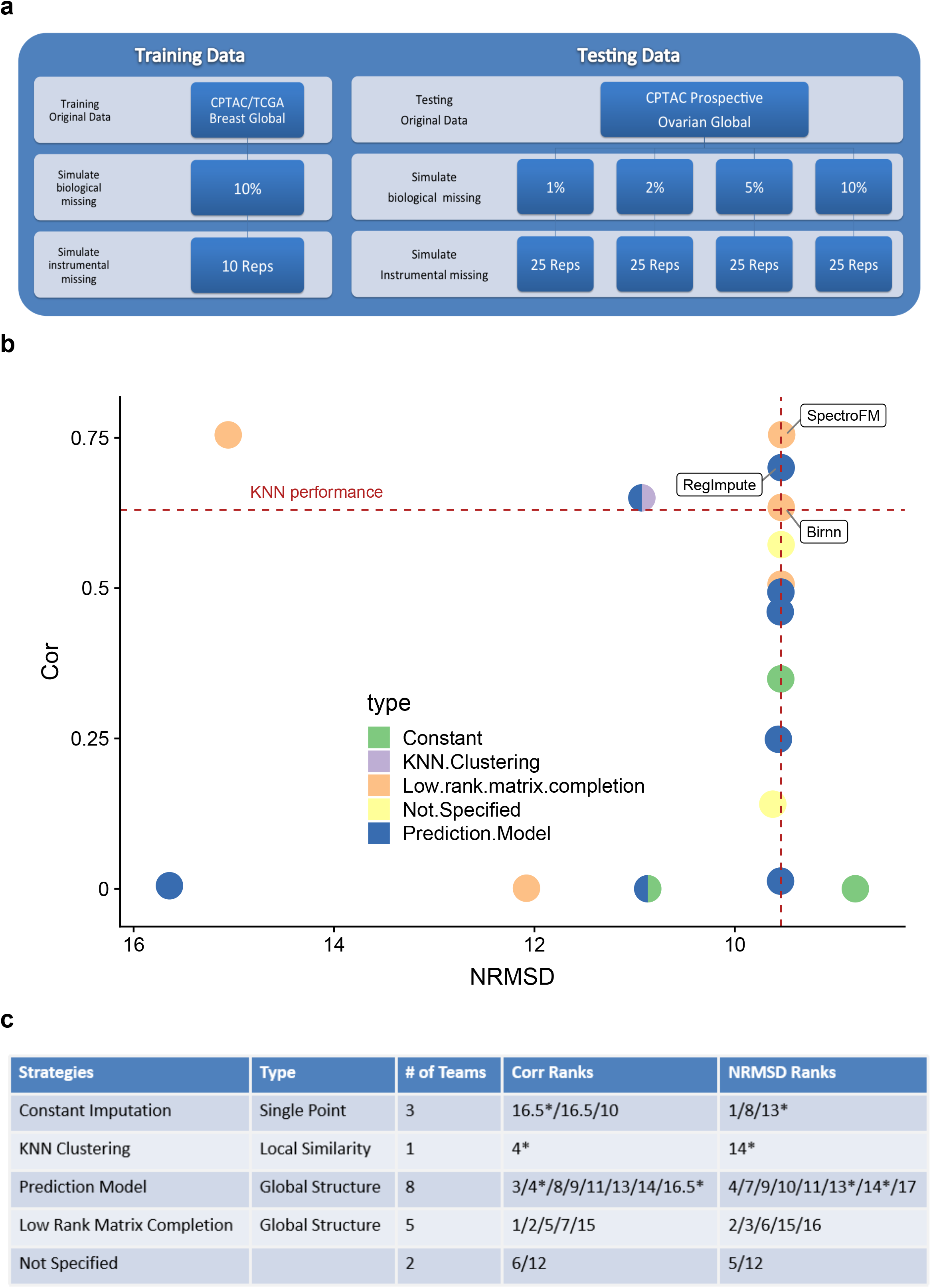
Proteomics Data Imputation Challenge competition design and performance results of participants. **(a)**: Design of data simulation in challenge phase. **(b)**: Cor and NRMSD evaluations of 17 submitted imputation algorithms. Different colors and shapes represent different imputation strategy categories. The dotted lines illustrate the performance level of KNN imputation. Three leading algorithms with better performance than KNN imputation have their names labeled. **(c)** Performance rank of all algorithms summarized for each strategy category (*algorithms using multiple strategies were listed multiple times in all relevant categories).

Imputation performances were assessed based on two metrics: protein-wise Pearson correlation coefficients (Cor) and normalized root mean squared deviation (NRMSD) between imputed and true values (see **Supplement A6**). Specifically, NRMSD was computed as the root of mean squared deviation normalized by the range of true values. The final ranking of participating teams during the challenge phase was determined by a tie breaking strategy applied on their performance evaluation (see **Supplement A6** and **Supplemental Table S2,S3**).

### The community phase

The goal of the community phase was to construct a consensus imputation algorithm by integrating multiple methods with diverse strategies. We not only utilized the winning algorithms from the challenging phase, but also leveraged existing tools that provide complementary strengths. We extensively evaluated the integration strategies and developed a baggingbased aggregation framework that enhances the robustness of the final algorithm. We refer to the final algorithm as DreamAI (Aggregated Imputation). Methodology and performance details of DreamAI are discussed in the *Result* Section.

To evaluate imputation performances, we utilized two replicate sets of protein abundance profiles of 32 tumors from the CPTAC/TCGA Ovarian Cancer Study (see **Supplement A2, A3** and **A5** for data set details and quality assessment). Specifically, the two replicate sets were independently generated by two proteomics labs: the Pacific Northwest National Laboratory (PNNL); and a proteomics lab from Johns Hopkins University (JHU). We thus referred to the two data sets of 32 samples as the PNNL-data and JHU-data respectively.

All methods were firstly applied to the PNNL-data (3027 proteins by 32 samples) to impute the missing values. The corresponding observed data points in the JHU-data were then used as approximation for the true values to evaluate the imputation results. There were 3700 missing values in the PNNL-data, and most (>99%) of them were observed in the JHU-data. To adjust for “background” abundance measurement differences between the two proteomics labs due to technical and/or biological factors, we employed the Scaled-Cor and NRMSD-**δ** for performance evaluation. Specifically, for each protein, background Cor and background NRMSD were first obtained using paired data points observed in both data sets. In addition, *imputation Cor* and *imputation NRMSD* were derived as the Cor and NRMSD between imputed values of the PNNL-data and the corresponding non-missing values in the JHU-data, respectively. Scaled-Cor of one protein was then calculated as the ratio of the imputation Cor to the background Cor of this protein. And NRMSD-**δ** was defined as the difference between the imputation NRMSD and the background NRMSD. In addition, to ensure robust evaluation, we restricted our performance evaluation on a subset of 289 proteins which had at least 5 missing data points and background Cor greater than 0.3.

### Validation of DreamAI performance on real application

To illustrate the impact of imputation on downstream data analysis of proteomics data, we applied DreamAI to a large TMT proteomics data set from a recent proteogenomics study of clear cell renal cell carcinoma (CCRCC) [35]. In this study, 103 treatment naïve renal cell carcinoma and 80 paired normal adjacent tumor (NAT) tissue samples were profiled using a proteogenomic approach wherein each tissue sample was homogenized via cryopulverization and aliquoted to facilitate genomic, transcriptomic, and proteomic analyses on the same tissue sample.

We calculated the concordance between RNAseq and global proteomics data by testing gene-wise Spearman correlations between these two types of data on the same set of samples. Evaluation of the concordance was based on proteins (n=2012) with at least one missing value in the tumor samples.

We also evaluated whether different treatments of missing values may impact statistical powers for detecting proteins associated with normal-tumor status or immune subtypes. We considered four data versions: the original abundance table with missing values, and the imputed abundance tables based on three different imputation methods: mean imputation, KNN imputation and DreamAI imputation. We focused on a subset of 92 genes in the CPTAC CCRCC proteomics data, whose imputed protein abundances by KNN and that by DreamAI are rather different (the NRMSD between the imputed abundance by KNN and that by DreamAI is greater than 0.2). We then performed Wilcoxon two-sample tests comparing tumor and NAT samples; and Kruskal–Wallis tests to screening for proteins associated with four different immune subtypes [35].

### Rational of Statistical Tests in the Study

The assumption of normal distribution is not always proper for proteomics abundance data from labeled mass spectrometry experiments. In **Figs S1e-f**, we illustrated the normality (Shapiro) test results for the CPTAC/TCGA Ovarian Cancer data sets. Majority of the proteins (>75%) in both the PNNL- and JHU-data showed significant deviation from the normal distribution (p < 0.05). Therefore, we chose to use non-parametric tests for all the statistical inference. Specifically, we employed the Spearman correlation based tests to assess RNA-protein concordance, the Wilcoxon Rank test to identify differentially expressed proteins between normal and tumor samples, and the Kruskal-Walis tests for screening for proteins associated with four different immune subtypes.

## Result

### Competition results in the challenging phase

We implemented two rounds of leader board competition and one final round of official competition evaluation during the challenging phase (see **Supplement A1**). Among 21 teams participating in this challenge, 17 received valid scores in the final round. Names and affiliations of all participants were listed in **Table S4**. The corresponding 17 imputation methods include 6 methods based on prediction models, 5 using matrix completion techniques, 2 relying on constant imputation, 2 employing multiple strategies and 2 other methods with missing algorithm strategies reports. The performances of these 17 algorithms were illustrated in **Fig. 2b, 2c**. Interestingly, diverse performances were observed for teams employing the same category of methods. For example, among the five low-rank matrix completion based imputation methods by five different teams, two showed superior performance, but the other three produced much less reliable results compared to the baseline imputation method, KNNimpute [17,18] (**Fig. 2b**). This observation suggests that customized treatment for labeled proteomics data in employing these imputation techniques is important to assure good performance. Also, as expected, the two methods based on constant imputation showed poor performances, suggesting this simple treatment does not work well for labeled proteomics data with complicated missing mechanisms.

The top three methods were SpectroFM, RegImpute, and Birnn (please see **Supplement A6** for ranking details). Both SpectroFM and Birnn used matrix completion techniques, while RegImpute employed prediction models. Methodology details are discussed in the following section and **Supplement A9**. The corresponding teams of the three winning algorithms --- SpectroFM, RegImpute, and Birnn --- were then invited to participate in the community phase.

### DreamAI: Methodology and Performance

In the community phase, we constructed a consensus imputation algorithm by integrating multiple methods with diverse strategies. The final product, DreamAI, is based on an aggregated imputation framework [36]. Specifically, it consists three major steps (**Fig. 3a**): generating 100 bagging sets with pseudo missing values based on the original data; imputing each bagging set with a consensus imputation strategy; and averaging imputed values of each missing spot across different bagging sets.

**Figure 3.**
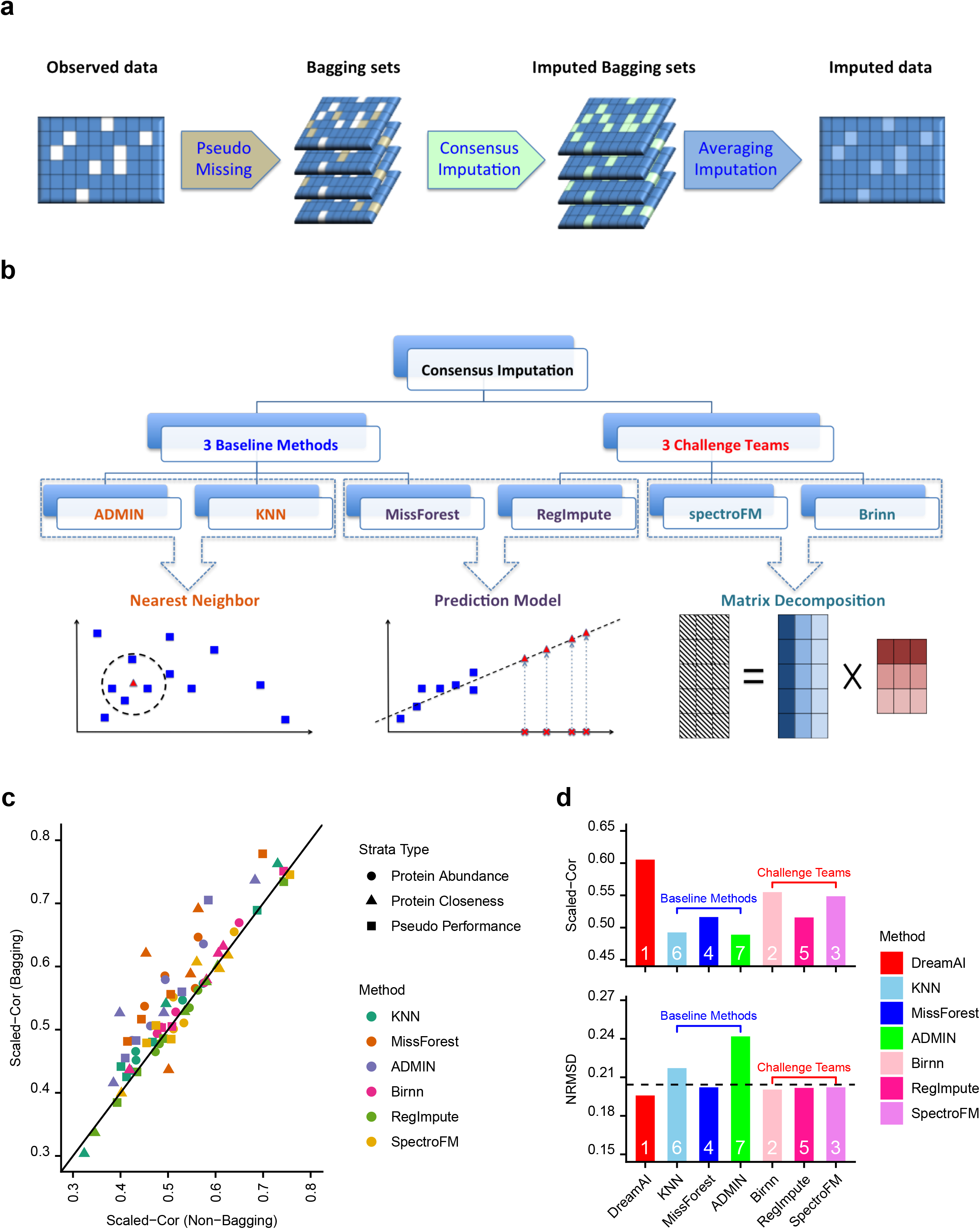
DreamAI algorithm and its performance. **(a)** Bagging procedure in DreamAI. Firstly, different set of pseudo missing are introduced to original observed data to generate a collection of bagging data sets. Then imputation is performed for each bagging set using the consensus imputation method. The final imputed matrix is the average of all bagging sets at the missing spots of the original data. **(b)** Consensus imputation method in DreamAI: average of 6 algorithms including 3 baseline methods and 3 winning algorithms from the Challenge. **(c)** Imputation performance (Scaled-Cor) of all individual imputation method with and without bagging strategy. Average Scaled-Cor are reported for different protein groups based on different protein closeness, average abundance, or pseudo missing performance evaluations. **(d)** Performance (Scaled-Cor and NRMSD) comparison between DreamAI and all individual methods. The dashed line in the NRMSD panel represents the background NRMSD between PNNL-data and JHU-data based on data points observed in both data sets. The white numbers labeled on the bars represent the ranks of the performance of each method.

#### The consensus imputation strategy

The central piece of DreamAI --- the consensus imputation strategy, is based on results from six imputation algorithms: the three winning algorithms in the challenging phase --- spectroFM: Team DMIS_PTG; RegImpute: Team Jeremy Jacobsen; Birnn: Team BruinGo; and 3 baseline algorithms --- ADMIN[6], KNN[17,18], and missForest[19] (**Fig. 3b**).

Both spectroFM and Birnn are based on low rank matrix completion methods. Specifically, spectroFM employs LibFM, a factorization machine library [37] to approximate the normalized protein abundance matrix (with missing values) with the product of two dense latent low rank matrices corresponding to proteins and samples respectively. In addition, a regularized Markov Chain Monte Carlo (MCMC) algorithm is implemented in spectroFM to solve the optimization problem. Birnn, which employs a similar low rank matrix decomposition framework, uses a different regularization technique --- the smoothly clipped absolute deviation (SCAD) penalty [38] --- to constrain the ranks of the decomposed matrices, and implements an iteratively reweighted nuclear norm (IRNN) [39] algorithm to solve the optimization problem (see **Supplement A9**).

Similar to missForest [19], RegImpute tackles the problem of imputation through prediction. The idea is to use observed abundances of other proteins (samples) to estimate the missing abundance of a given protein (sample). Specifically, RegImpute utilizes ridge regressions [40], and incorporates an iterative procedure to refit the prediction models leveraging the imputed values from the last iteration (see **Supplement A9**). This iterative procedure helps improve the prediction accuracy, and usually converges after 10 iterations.

KNN based imputation, the most commonly used imputation strategy in omics studies, can also be viewed as a prediction approach: a small set of features (samples) in the neighborhood of the feature (sample) to be imputed are used to fit a prediction model, which often takes the form of a linear combination (weighted average). ADMIN [6] can be viewed as an enhanced version of KNN. It specifically models the abundance-dependent missing mechanism in labeled proteomics data sets, and uses the joint likelihood of protein abundances and missing mechanisms to calculate the optimal weights for predicting the missing values (see **Supplement A8**).

In addition, when selecting baseline methods to be included in DreamAI aggregation, we also considered *pcaMethods* [30], a low-rank matrix completion method that has been applied to missing value imputation of labeled proteomics data [31]. However, the performance of *pcaMethods* is substantially worse than that of KNN, MissForest, and ADMIN on the CPTAC/TCGA Ovarian Cancer data set (**Fig S3**). Thus, we did not include this algorithm in the final consensus of DreamAI.

All selected methods provide complementary strengths. While the low rank matrix completion based methods effectively leverage the strong global covariance structure among proteins, the prediction-based methods provide more flexible imputation solution based on small neighbors (individual features) in the data. In addition, missForest helps to capture non-linear relationship among proteins, and ADMIN directly utilizes the abundance-dependent missing trend in proteomics data. Thus, by aggregating all these strategies in an effective way, we expect to achieve more accurate and robust imputation performance. Specifically, we propose to average the imputation results of all the 6 methods on one data set as the consensus imputation strategy. The bagging procedure, described below, makes this simple average rather robust and effective.

#### Model aggregation through bagging

A modified bagging strategy is adopted in DreamAI to improve the robustness and accuracy of imputation algorithms. Instead of sub-sampling subjects or proteins, DreamAI generates “bagging” (perturbed) data matrices by setting a small subset of observed data points in the original data matrix as pseudo NAs. Specifically, these data points were selected according to a probability model reflecting the abundance-dependent missing mechanism with parameters estimated based on the original data matrix. Then DreamAI applies imputation algorithms on a collection of bagging matrices with both true and pseudo missing values, and reports the average of the imputed values of each missing spot across all bagging matrices as the final imputed values.

To illustrate the benefit of bagging aggregation on imputation, we applied individual imputation method with or without bagging aggregation on the PNNL data. Specifically, we utilized 100 bagging matrices, and set the missing rates in the bagging matrices to double that of the original data set. For each method, Scaled-Cor between imputed values and the observed “true” values from the JHU data set of the corresponding data points were used for evaluation (see *The Community Phase* section for definition of Scaled-Cor). **Fig 3c** illustrates Scaled-Cor of imputation results from methods using (y-axis) and not using the bagging aggregation strategy (x-axis) across protein groups with different characteristics. Specifically, protein groups were defined according to **protein closeness scores, pseudo missing performances**, or average **protein abundances**. For a given protein, the protein closeness score refers to the average correlation between this protein and its 50 closest neighbors; the pseudo missing performances were calculated as the NRMSD between the imputed values of the pseudo missing data points introduced during the bagging procedure and their “true” abundances; and the average protein abundances were derived based on observed protein abundance measurements in the PNNL-data. More details of definitions of these protein groups were provided in **Supplement A7**.

As shown in **Fig. 3c**, the results based on bagging aggregation showed overall improved Scaled-Cor compared to those without using bagging aggregation.

Additionally, the improvement from bagging is more dramatic for baseline methods than the winning algorithms from the challenging phase.

#### Overall performance evaluation

We compared the performance of DreamAI with that of each individual imputation algorithm (with bagging). The average Scaled-Cor and NRMSD based on all proteins are shown in **Fig. 3d**. DreamAI achieves higher Scaled-Cor and lower NRMSD than all the six individual imputation methods. Specifically, the imputation accuracy of DreamAI, as measured by Scaled-Cor, is about 20% greater than KNN and ADMIN, and 15% greater than missForest. In addition, the performance of DreamAI was also compared to that of *pcaMethods,* and a 50% improvement on Scaled-Cor was observed (**Fig S3**). In addition, the dashed line in the NRMSD plot represents the reference NRMSD based on all paired data points observed in both the PNNL and the JHU data sets. Interestingly, NRMSD of DreamAI is smaller than the reference NRMSD, implying superior performance of DreamAI.

As illustrate in **Fig. 3d**, the three winning algorithms from the Challenge all outperformed the three baseline methods, which is consistent our observations from the challenge phase. An immediate question, then, is whether it helps, in the aggregation exercise, to include any or all of the baseline methods, which have suboptimal performances. We thus also evaluated strategies of aggregating none or a subset of the baseline methods in DreamAI. As illustrated in **Fig. S2a**, without any of the baseline methods, the Scaled-Cor of imputation result is about 13% lower than the result from aggregating all 6 methods. This clearly demonstrates the benefit of aggregating methods with complementary strengths. Moreover, ADMIN appears to be a more important player than KNN and missForest, such that the Scaled-Cor drops more if ADMIN were ommitted from the aggregation, compared to when missForest or KNN were left out. This illustrates the benefit of incorporating the abundance dependent missing mechanism, a common feature of proteomics data, in the imputation framework. Between KNN and missForest, KNN is less helpful in the aggregation, such that the method by leaving KNN out achieves even slightly better performance in terms of Scaled-Cor. More detailed investigation further suggests that KNN helps only for proteins with close neighbors and high abundances (**Fig. S2b-c**).

In practice, the DreamAI R-package provides the flexibility for users to specify any combination of the 6 individual methods to perform DreamAI imputation. When the data dimension or computational cost is not a concern, one may choose to include ADMIN and missForest, in addition to the three winning algorithms, to achieve the optimal performance. When the data matrix has a large dimension, computational time required by missForest could be substantial, and the users may choose to include ADMIN and KNN instead of missForest to balance the tradeoff between performance and computational burden.

#### Performance across different protein groups

To further understand the impact of various protein characteristics on the imputation results, we assess the performance of DreamAI among protein groups with different **protein closeness scores, pseudo missing performances**, or average protein abundances as defined in the previous section and in the **Supplement A7**.

As illustrated in **Fig. 4a**, imputation performance of DreamAI, in term of scaled-Cor, shows an increasing trend with protein closeness. Moreover, the improvement of DreamAI over KNN is the most dramatic, more than 65%, for the protein group with the lowest closeness. This suggests the advantage of leveraging the information in the whole data set for data points with uninformative neighbors when performing imputation. A similar pattern is observed based on NRMSD-**δ** as well (**Fig. 4a**).

**Figure 4.**
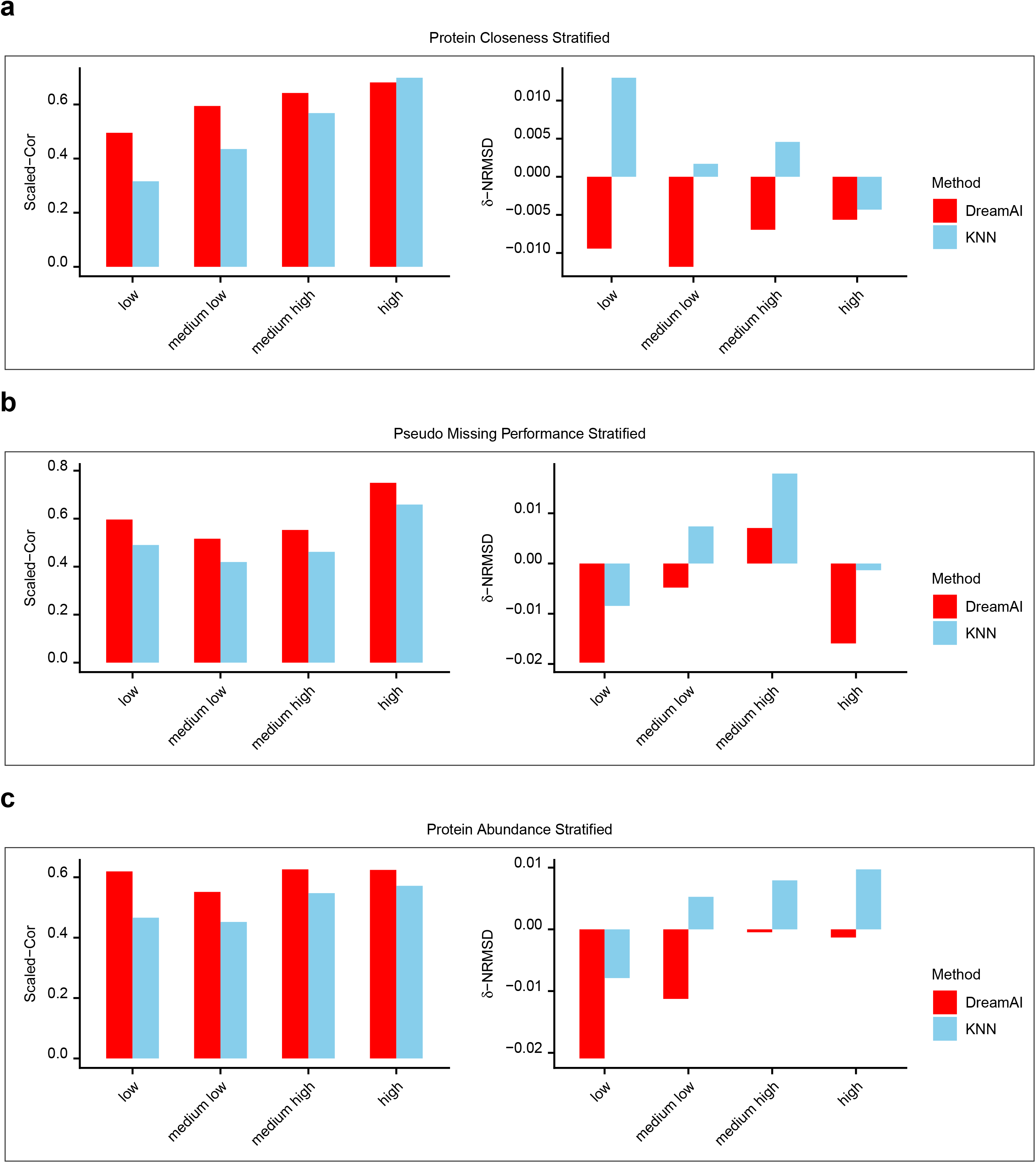
Scaled-Cor (left) and NRMSD-δ (right) of DreamAI and KNN for different protein groups (x-axis) based on the CPTAC/TCGA ovarian data. The protein groups are derived based on **(a)** protein closeness; **(b)** pseudo missing performance; and **(c)** average protein abundances. For one protein, the protein closeness score refers to the average correlation between this protein and its 50 closest neighbors; the pseudo missing performances were calculated as the NRMSD between the imputed values of the pseudo missing data points introduced during the bagging procedure and their “true” abundances; and the average protein abundances were derived based on observed protein abundance measurements in the PNNL-data. More details of definitions of these protein groups were provided in **Supplement A7**.

Across the four protein groups with different pseudo missing performance evaluations, both DreamAI and KNN showed better imputation accuracy in term of Scaled-Cor for the group with the best pseudo missing performance than the others. The improvements of DreamAI over KNN, however, are quite comparable across the four clusters (**Fig. 4b**).

While protein abundance correlates with the imputation performance of KNN, it does not show obvious association with performance of DreamAI (**Fig. 4c**). And DreamAI showed the biggest improvement over KNN for the protein group with the lowest abundances. NRMSD-**δ** of both DreamAI and KNN appeared to be negatively associated with the protein abundance, which seems to imply that NRMSD depends on the scale of the value to be imputed, and thus its interpretation needs to be taken with cautious.

### Imputation helps gain biological insights

To illustrate the benefit of proper imputation of proteomics data on downstream proteogenomic analysis, we applied DreamAI on the global proteomics data set of 103 treatment naïve renal cell carcinoma and 80 paired normal adjacent tumor (NAT) tissue samples from the CPTAC CCRCC study [35]. The data set contained protein abundance measurements of 9209 genes that were detected in at least 50% of the samples. 2059 of the 9209 genes had missing measurements in at least one sample. The overall missing rate of the protein abundance matrix of these 2059 genes was 20.4%, and sample wise missing rate ranges from 2.5% to 7%. The abundance dependent missing (MNAR) trends in the proteomics data of tumor and NAT samples are illustrated in **Fig. 5a** and **S4a** respectively.

**Figure 5.**
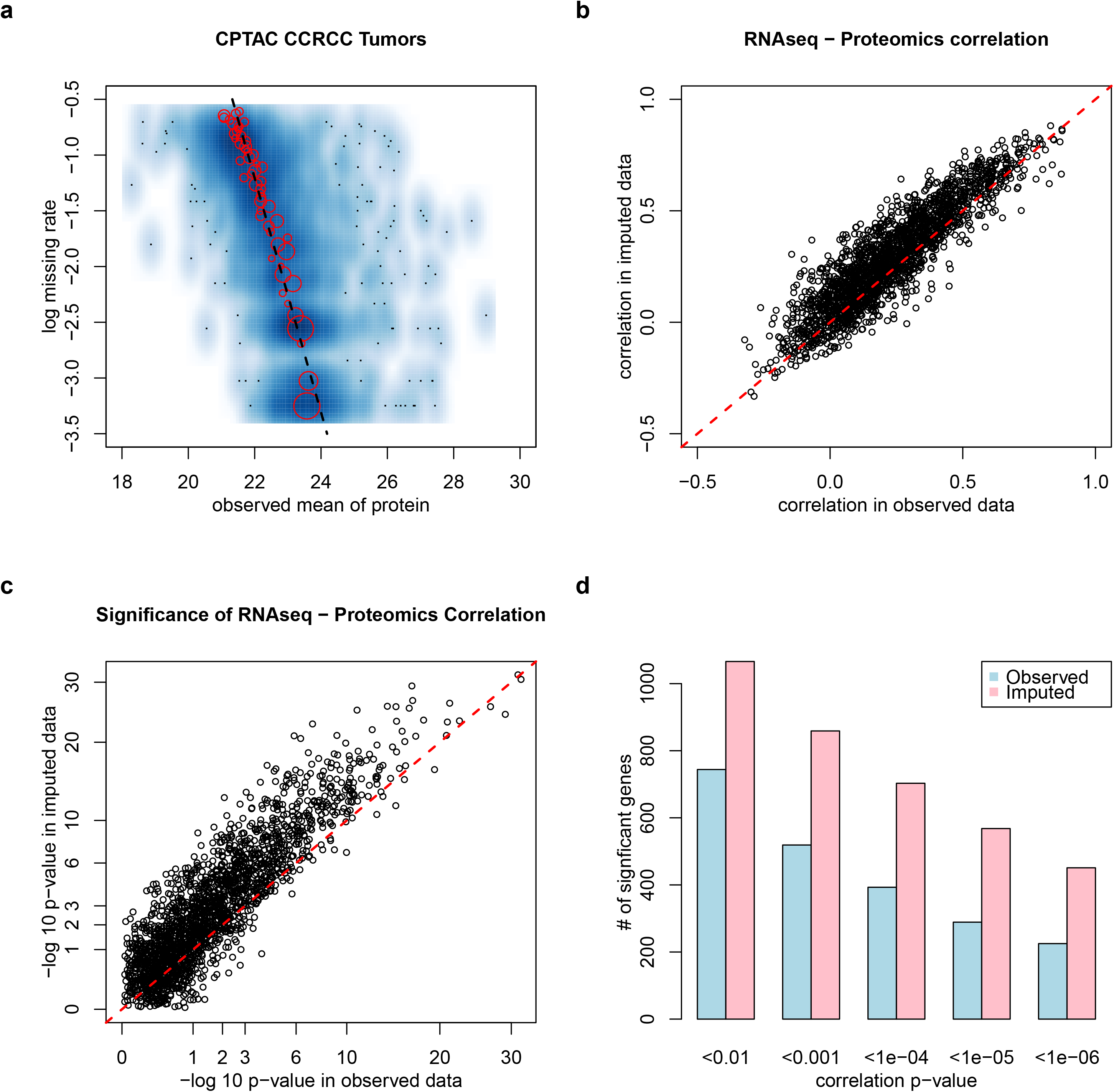
For a set of CPTAC CCRCC tumors, proteomics data with DreamAI imputation shows improved concordance with their corresponding transcriptome data. All “correlation” labels refer to the Spearman correlation in this figure. **(a)** Scatter plot of protein-level missing rates vs. mean protein abundances based on observed values in the global proteomics data of 103 CCRCC tumor samples [35]. **(b)** Scatter plot of protein-RNA association (Spearman correlation) between RNA and proteomics data with DreamAI imputation (y-axis) vs. the association based on non-imputed data (x-axis). **(c)** Scatter plot of significance (− log 10 p-value) for Spearman correlation between RNA and proteomics data with DreamAI imputation (y-axis) vs. that for the data without imputation (x-axis). **(d)** Number of genes showing significant protein-RNA Spearman correlation based on proteomics data with DreamAI imputation (pink) or without imputation (blue) at different p-value cutoffs.

We first considered the 103 tumor samples and assess the protein-RNA concordance for the protein abundance matrices with and without DreamAI imputation. Since the noises from any analytical, experimental or data preprocessing steps in the proteomics data should be independent from that in the RNA data, better protein-RNA concordance shall reflect richer information that is biological relevant in the corresponding data sets. As illustrated in **Figure 5b**, for 2012 proteins with at least one missing value in tumor samples, their gene-wise protein-RNA Spearman correlations based on DreamAI output were significantly higher than that without imputation (pvalue<10e-16). The DreamAI result also led to a greater number of genes with significantly non-zero Spearman correlation at various p-value cutoffs (**Figs 5c, 5d**). Parallel analysis applied to proteogenomics data of NAT samples reveals similar improvement of protein-RNA concordance based on protein abundance table with DreamAI imputation over that without imputation (**Fig. S4**).

We also observed increased power of biological signal identification based on DreamAI imputation results, as compared to other imputation strategies. Specifically, **Fig. S5a** illustrated the p-value distributions of the Wilcoxon two-sample tests comparing tumor and NAT samples based on either the original abundance table or those resulted from different imputation strategies. The p-values based on the DreamAI imputation tended to be more significant than those resulting from the mean- or KNN-imputation, as well as that of no imputation. Moreover, **Fig. S5b** shows a similar benefit on power for screening for proteins associated with four different immune subtypes [35] by using DreamAI imputation over other imputation strategies.

Furthermore, to investigate whether imputation may induce (artificial) changes of the abundance distributions, we assessed whether/how different imputation strategies could impact the standard deviations (SD) of individual protein abundances (**Figs. S5c**). First, as expected, for a large number of proteins, the mean-imputation resulted in decreased SDs, which might cause artificially inflated false discovery rate in downstream statistical inferences. The KNN imputation leads to both deflated and inflated SDs for different groups of proteins. DreamAI, on the other hand, demonstrated the least perturbation on the SDs of protein abundances.

These examples illustrate the advantage of using proteomics data with DreamAI imputation in downstream statistical analysis over other alternative strategies. Nevertheless, we also want to note that without knowing the ground truth in the real data, one needs to interpret the power comparison for differential expression analysis with caution.

### Software and Computational Time

An R package of DreamAI has been implemented and is available to public at Github (https://github.com/WangLab-MSSM/DreamAI). We have also provided a Docker image of the software at https://hub.docker.com/r/cptacdream/sub1.

To illustrate the computational time of the algorithm, we applied the docker version of DreamAI to the CPTAC CCRCC proteomics data (9209 genes/proteins by 207 samples) using a cloud based "medium" size Linux virtual machine comprised of 2 vCPUs and 4gb RAM with an Ubuntu operating system. The imputation took 4hr50min when all 6 imputation methods included in the DreamAI package were used for the ensemble. The running time reduced to 1hr50min when missForest is not included in the ensemble procedure. Thus, when computational burden is a concern, the users could choose to omit missForest in the DreamAI function to speed up the imputation. This strategy, in general, will not sacrifice much of the imputation accuracy as illustrated in the CPTAC/TCGA ovarian cancer data example (Figure S2). Please see more discussion on the recommended choices of collection of ensemble methods in DreamAI in the ***Overall performance evaluation*** section.

Here we want to note that the computational time of DreamAI on a typical proteomics data set is rather comparable to other computational steps in a proteomics data analysis pipeline. Protein identification and quantification often takes much longer time than the imputation procedure. For example, it takes about 15hrs to produce a protein level abundance matrix from raw file of the CPTAC CCRCC proteomics data on a horizontal server with 56 cores (Intel(R) Xeon(R) CPU E5-2690 v4 @ 2.60GHz), and 500 RAM by computational tools including: MSFragger and Philosopher[35,42,43].

## Discussion

How to handle missing values in MS based proteomics data has been a long-standing challenge in proteomics research. As the study size increases, the issue of missing values becomes more significant, as data from more mass spectrometry experiments need to be merged together. On one hand, the isobaric labeling technique greatly enhances the quantitation precision and experiment throughput; on the other hand, it further exacerbates the missing data problem. With experimental induced multiplex-level missing pattern as well as the abundance dependent missing trend, proteomics data from labeled MS experiments cannot be properly or effectively analyzed by using observed data only (either ignoring all features with missing samples or ignoring subsets of samples with missing data points in feature-wise modeling).

Another strategy to handle missing data is through imputation, which has been widely adopted in many research fields, such as image processing, single-cell RNAseq studies, and label free proteomics data analysis. Its usage in proteomics data from labeled MS experiments is still limited, largely due to a lack of a benchmark imputation method suitable for this type of data. Because of the complicated missing structure in labeled proteomics data, imputation tools developed for other data types do not apply or do not perform well.

The goal of this study is to develop a benchmark imputation algorithm for labeled proteomics data sets. Specifically, we conducted the NCI-CPTAC DREAM Proteogenomics Imputation Challenge to achieve this goal through crowd learning. 21 teams from a broad range of research fields participated in the Challenge and contributed diverse expertise. As expected, many general imputation algorithms used in other disciplines/applications did not perform well on labeled proteomics data sets. Indeed, only a subset of teams achieved better performance than the KNN imputation on Challenge data sets, suggesting customization of the imputation algorithm for labeled proteomics data is important in order to effectively tackle this problem.

The three winning teams from the Challenge further participated in a collaborative phase, and we jointly developed the final algorithm --- DreamAI --- an ensemble based imputation method. DreamAI employs a bagging framework to aggregate results from 6 diverse imputation methods: three winning algorithms from the Challenge (two based on low-rank matrix completion and one based on prediction model fitting), as well as three baseline imputation methods which have been used in previous proteogenomics data analysis --- KNN, ADMIN, and missForest [5,6,20,21]. This ensemble strategy of DreamAI leads to greatly improved performance compared to that of individual algorithms. The imputation accuracy on Scaled-Cor of DreamAI is 15-50% better than that of the individual baseline tool, or 9-15% better than that of the individual winning algorithm on the CPTAC/TCGA Ovarian Cancer proteomics data set.

The bagging framework in DreamAI not only enhances the imputation performance, but also helps one gain insights on the imputation quality of each feature. Specifically, for a given feature, DreamAI estimates its imputation quality using the Cor between the true and imputed values of pseudo missing data points of this feature across different bagging iterations. In the CPTAC/TCGA Ovarian Cancer data application, the Cor assessment for the protein group with the best pseudo missing performance is 0.75, at least 26% higher than the remaining protein groups. Therefore, the pseudo missing performance score of each feature is informative to shed light on feature-specific imputation quality.

Since imputation is an unsupervised learning problem, it has been a challenging task to objectively assess the performance of imputation methods. Thus, one of the major efforts during the Dream Challenge was to create high-quality benchmark simulation data sets to objectively evaluate imputation performances. Specifically, simulations were set up to mimic missing patterns in real proteomics data sets as closely as possible. Multiple training and testing data sets with varying proportions of biological and experimental missing rates, as well as different degrees of abundance dependent missing trend were generated based on global proteomics data from CPTAC breast and ovarian cancer studies [1,7]. Especially, global proteomics data from the CPTAC Prospective Ovarian Cancer Study [7], which were not publicly available during the Challenge Phase, were used to generate the testing data sets.

Moreover, to complement the usage of simulated data sets during the Challenge phase, in the community phase, we utilized the CPTAC/TCGA Ovarian Cancer proteomics data set [2], which contained proteomics profiles of two replicate biological samples of 32 ovarian tumors. This provided a unique opportunity to directly assess imputation performances on real missing data points in cancer proteomics studies.

Although we provided NRMSD values on all examples, we used Cor as the main metric to evaluate imputation performance. NRMSD measures the distance between the imputed values and the true values of missing data points normalized by the varying range of abundances of each protein. Despite being a normalized distance measure, NRMSD still depends on the scale and distribution of the protein abundances. On the other hand, Cor is a scale free measure. As illustrated in **Fig. 4c**, among protein groups with different mean abundance levels, performance based on Cor is very stable, while NRMSD has an obvious trend to be positively associated with protein mean abundances.

The benefit of using imputed data in downstream analyses stems from the improvement of sample size and thus the analysis power. As illustrated in the application on CPTAC CCRCC proteomics data, imputation helps to improve the overall RNA-protein concordance. Note, as reported by multiple proteogenomic cancer studies [1,7,8,35,44], only 50%-75% genes show significant Spearman correlations between their RNA expression and protein abundances, as a large number of proteins are subjected to post-translational modifications. For example, in CPTAC CCRCC tumor and NAT data, about 74% and 52% of cis mRNA-protein pairs showed significant positive Spearman correlations respectively [35]. On the other hand, it’s reasonable to assume that noises coming from any analytical, experimental or data preprocessing steps in the proteomics data should be independent from that in the RNA data. Thus, for the protein data set showing higher levels of concordance with the corresponding RNA data, it is more sensible to assume the data set bears more relevant biological information, than to assume some artificial effects in the data set contributes to the better mRNA-protein concordance by chance.

In the example of the CPTAC CCRCC study, using imputation result by DreamAI also leads to more significant p-values than that of other alternative strategies, when screening for proteins associated with tumor/normal status or immune subtypes. A further investigation on the impact of imputation on the SD of individual protein abundances suggests that DreamAI has the least perturbation on the SDs of protein abundances. Nevertheless, we want to note that without knowing the ground truth in the real data, one needs to interpret the power comparison for differential expression analysis with caution.

### Other considerations in handling labeled proteomics data

In the labeled MS experiments, it is recommended to include a common reference sample in each multiplex of the same study. Then, by using the relative abundance to the reference sample in each multiplex for downstream analysis, one can effectively remove the batch effect induced by different experimental variations across multiplex runs. As to the choice of the reference sample, it is preferred to use one as similar as possible to the target samples in the study. For example, in the CPTAC ovarian cancer and CCRCC projects, the reference samples were created by pooling equal amounts of individual tumor samples to be profiled in the studies.

The usage of a common reference sample in each multiplex in isobaric labeled proteomics experiments also helps with the peptide-protein roll-up during protein quantification. Specifically, rolling up from peptides to proteins can be performed at the log-ratio intensity level (i.e. log-ratio between intensity of a target sample and intensity of the reference sample in the same iTRAQ/TMT multiplex for one peptide). This strategy greatly improves the robustness and precision of protein quantification, while at the same time, effectively reduces the missing data percentage in protein level data compared to the peptide-level data. Thus, for isobaric labeled global proteomics experiments, it is preferable to perform biological/clinical data analysis based on the protein level data. And consequently, we would recommend performing imputation on the protein/gene level data as well. On the other hand, for data analysis of label free proteomics data, it has been suggested that directly model peptide abundance could be more efficient than performing imputation at the protein abundance level [12]. This is because the summary (or average) based peptide-protein intensity roll-up used for label free proteomics data is vulnerable to many confounding factors. Modeling the peptide level abundances directly could effectively get around the variabilities induced in the roll-up step.

For phospho-proteomics experiments, since phospho-site is the biological unit for downstream analysis, it is more meaningful to work with the quantification at phospho-site level and perform imputation on phospho-site level data directly. Intuitively, since experimental noise in the data, including sample loading variations and batch effects could result in misleading patterns during imputation, we recommend performing imputation on data sets after proper pre-processing treatments, such as normalization and batch correction. In general, if the study contains samples with different disease statuses or from different treatment groups, it is better to have a randomized design to allocate samples from different groups evenly across multiplexes. Otherwise, batch correction will be needed in addition to the adjustment using the common reference samples. For example, in the CPTAC CCRCC proteomics experiment, batch effect due to TMT multiplex was confounded with the tumor-normal status due to imbalanced experimental design for a small subset of TMT multiplexes. Therefore, a batch correction step was adopted before imputation and downstream analysis were carried out. [35]

In the real data analysis, we removed features with missing rate higher than 50% before imputation and downstream analysis. The choice of 50% cutoff is a trade-off between imputation accuracy and information (data feature) loss of the data set. For features with a high missing rate, the tasks to accurately identify close neighbors or to fit prediction models based on observed data points become very challenging due to the sample size limitation. In a previous work, the authors suggested that, in general, imputation methods perform better on features with less than 50% missing values than on features with more than 50% missing values [12]. Additionally, in downstream analyses, it is preferred that the observed data points outweigh the imputed data points to ensure robustness. Thus, we settled with a cutoff at 50%.

Although DreamAI has a general framework and can be applied to other proteomics data from label free experiments, its performance on those applications warrants future study. In addition, for proteomics data from targeted mass spectrometry experiments, such as MRM (multiple reaction monitoring), imputation could be less of a concern due to the relatively low missing rate. However, MRM experiments right now can handle at most a few hundred proteins/peptides in one run, and thus are not suitable for deep profiling in discovery studies.

Application of SVM (support vector machine) or ANN (artificial neural network) on proteomics data imputation were not used by any of the 17 teams during the challenging, nor reported in any literature to the best of our knowledge. However, ANN based methods, which can be viewed as a non-linear extension of Low Rank Matrix Completion, have been applied to solve imputation and batch correction for single cell RNAseq data recently [41]. Machine learning methods including ANN could be presumably useful for proteomics data imputation, which warrants future investigations.

## Abbreviations

iTRAQ: isobaric tags for absolute and relative quantification
TMT: tandem mass tags
PTM: post-translational modifications
MNAR: missing not at random
KNN: K nearest neighbors
PCA: principal component analysis
CPTAC: Clinical Proteomic Tumor Analysis Consortium
TCGA: The Cancer Genome Atlas
CCRCC: clear cell renal cell carcinoma
Cor: Pearson Correlation Coefficient
NRMSD: normalized root mean squared deviation
PNNL: Pacific Northwest National Laboratory
JHU: Johns Hopkins University
NAT: normal adjacent tumor
MCMC: Markov Chain Monte Carlo
SVM: support vector machine
ANN: artificial neural network

## ACKNOWLEDGEMENT

We would like to thank the National Cancer Institute’s Clinical Proteomic Tumor Analysis Consortium (CPTAC), a comprehensive and coordinated effort to accelerate the understanding of the molecular basis of cancer through the application of proteogenomics, on providing the data used in this challenge and making it freely available to the public. We also like to thank Dream Challenges organization for providing the opportunity to encourage researchers all around the world to take part in this cutting-edge research topic and all the participants in this challenge for building the algorithms and submitting the results. This work was partly supported by grants U24 CA210993 and U24 CA210972 from the National Cancer Institute Clinical Proteomic Tumor Analysis Consortium (CPTAC).

## COMPETING FINANCIAL INTERESTS

The authors declare no competing interests.

## Main Figure Legends

## Notes

### Competing Interest Statement

The authors have declared no competing interest.

### Summary of Updates

We added a new section to describe the data sets and related bioinformatic and statistical methods used in the challenge and the follow up study. We have improved the representation of the manuscript and provided more technical detail in the new method section and supplements. Author affiliations and supplemental files were also updated.

