## supplemental materials for "DreamAI: algorithm for the imputation of proteomics data"

### Supplement A: Supplemental Methods

#### A1. Challenge Design

To help the Challenge participants to train their models, we implemented two rounds of leader board competition and one round for final competition evaluation during the challenging phase. Different training/testing data sets were used at each of the leader board or final rounds. During the leader board rounds, the participants were allowed to submit their algorithms multiple times, and the testing result of each submission was published. The participants could further tune their algorithms based on the feedback from the leader board rounds. In the final round, only one submission of each team was permitted to assure fair competition. Final performance and ranking of all participating teams were summarized and released after the closing of final submission.

**A2. Data Sets Summary**

Data sets used in the Challenge included both published and unpublished data (during the challenging period) from CPTAC/TCGA breast and ovarian cancer studies (**Table S1**). Global proteomics data from CPTAC/TCGA breast cancer study (iTRAQ) [1] was used to generate training data sets during the challenge phase. Global proteomics data from the CPTAC Prospective Ovarian Cancer Study (TMT) [2], which were not publicly available during the Challenge Phase, were used to generate testing data sets. Note, we chose to use different real proteomics data of different tumor types and experimental platforms to generate the training and testing data sets, because this helps to assure fair evaluation of the performance of the imputation algorithms.

Global proteomics data from the CPTAC/TCGA Ovarian Cancer Study [3] were used for imputation performance evaluation in the Community Phase. In the manuscript, we further utilized phosphoproteomics data sets from CPTAC/TCGA and Prospective Ovarian Cancer Studies [2,3] to illustrate missing rates and missing patterns in labeled proteomics data sets. Moreover, global proteomics data set from CPTAC CCRCC Discovery Study [4] was used to evaluate the performance of DreamAI compared to other missing value handling strategies in the manuscript.

For all aforementioned data sets, MS1 intensity (peak area) based quantification was used for protein abundances, which were further elaborated in **Section A.2**. Other details of these data sets were provided in **Table S1**.

**Table S1.** Summary of Data Sets used in the Challenge and the manuscript.

| Cohort | # Protein | # Sample | Plat- form | Usage | Publically Available* | PUBMED ID (Year) |
| --- | --- | --- | --- | --- | --- | --- |
| CPTAC/TCGA Breast Cancer | 11585 | 80 | iTRAQ (global) | Challenge Phase ---Training | Yes | 27251275  (2016) |
| CPTAC/TCGA Ovarian Cancer (JHU) | 5122 | 122 | iTRAQ (global) | Community Phase | Yes | 27372738  (2016) |
| CPTAC/TCGA Ovarian Cancer (PNNL) | 3080 | 84 | iTRAQ (global) | Community Phase | Yes | 27372738  (2016) |
| CPTAC/TCGA Ovarian Cancer Phospho (PNNL) | 12434 phospho-site | 69 | iTRAQ (phospho) | Manuscript---Missing rate/pattern illustration | Yes | 27372738  (2016) |
| CPTAC Prospective Ovarian Cancer | 10853 | 103 | TMT (global) | Challenge Phase ---Testing  Manuscript---Missing rate/pattern illustration | No | 32529193  (2020) |
| CPTAC Prospective Ovarian Cancer Phospho | 40966 phospho-site | 103 | TMT (phosphor) | Manuscript---Missing rate/pattern illustration | No | 32529193  (2020) |
| CPTAC CCRCC Discovery | 11357 | 183 | TMT (global) | Manuscript---Imputation Evaluation | No | 29988108  (2018) |

**Yes/No indicates whether the dataset was publicly available during the Challenging phase.*

**A3. Download and Preprocessing of the Protein Abundance Matrices**

***CPTAC/TCGA Breast Cancer Study***

Raw data files of iTRAQ LC-MS/MS experiments were downloaded from the CPTAC public data portal (https://cptac-data-portal.georgetown.edu/). Protein identification and quantification were performed using Spectrum Mill software package v5.0 with RefSeq database as described in the study publication [1].

For each iTRAQ multiplex experiment, the output Peptide-Spectrum Match (PSM) table from Spectrum Mill reported the mapped peptide sequences, related protein and gene names, MS1 parent ion intensities and MS2 isobaric label intensities of each spectrum detected in the MS2 step. Here, the MS2 intensities of isobaric labels reflect the abundance proportions of the peptide in individual samples of the multiplex.

In this study, each iTRAQ multiplex contained three tumors plus a fourth common internal reference sample. The reference sample comprised 10 individual tumors of each of the 4 major breast cancer intrinsic subtypes and served as an internal standard for all proteins and phosphoproteins quantified in this study. Relative abundances of proteins were determined in Spectrum Mill using iTRAQ reporter ion intensity ratios (tumor samples versus the reference sample) from each PSM. A protein-level or phosphosite-level iTRAQ ratio was calculated as the median of all PSM level ratios contributing to a protein subgroup after proper quality control [1].

MS1 parent ion intensities were summarized at protein level for each iTRAQ multiplex from the PSM table. The MS1 intensity of the multiplex represented the total abundance of all samples in the multiplex, and was further broken down to sample level summary with the proportion determined by MS2 ratios. Consequently MS1 intensities on gene level were produced for all samples in the experiment. In the end, sample-wise normalization was performed by shifting the median of each sample to the median of entire protein abundance matrix; and batch effects due to iTRAQ multiplex were removed using the mixEMM algorithm [5] based on the common reference samples in each multiplex.

***CPTAC Ovarian Cancer Studies***

Raw data files of CPTAC/TCGA Ovarian Cancer Study(iTRAQ)[3] and CPTAC Prospective Ovarian Cancer Study(TMT)[2] were downloaded from the CPTAC public data portal (https://cptac-data-portal.georgetown.edu/). To be consistent with the other 2 sub-challenges [6] in this Dream Challenge event, Common Data Analysis Pipeline (CDAP) [7] was used for database search and quantification on these data sets.

In these studies, each iTRAQ or TMT multiplex contained a common internal reference sample. The reference samples were made through pooling equal amounts of individual tumor samples in each study, and served as internal standards for all proteins and phosphoproteins quantified in these studies.

For each iTRAQ/TMT multiplex experiment, the output Peptide-Spectrum Match (PSM) table from CDAP reported MS1 parent ion intensities and MS2 isobaric label intensities, which were then used to re-scale the MS1 parent ion intensity of the peptide to produce the sample level peptide abundance measurements. To roll up peptide abundances to the protein level, intensities of peptides mapping to the same proteins were averaged and log2 transformed. Note, peptides mapped to multiple protein/genes and peptides with substantial missing values (over 50% across all samples) were excluded in the roll-up. Moreover, only protein/genes with at least two unique peptides were included in the protein abundance table. In the end, sample-wise normalization were performed by shifting the median of each sample to the median of entire protein abundance matrix; and batch effects due to iTRAQ/TMT multiplex were removed using the mixEMM algorithm [5] based on the common reference samples in each multiplex. For preprocessing of the phosphoproteomics data of the Ovarian Cancer studies, similar procedure was taken.

Missing data summary and abundance dependent missing patterns were illustrated with the data from the above pipeline. Beside the standardized pipeline, we also generated a version of the global proteomics data of the CPTAC Prospective Ovarian Cancer Study using relaxed filtering on the peptide-protein/gene roll-up (no requirement of unique peptides). This resulted in a global proteomics data with larger dimension, so we could generate the testing data sets of the similar size as the training data sets.

***CPTAC CCRCC Discovery Study***

We utilized the (MS1 intensity based) protein abundance table associated with the study publication [4]. Specifically, raw data files of TMT LC-MS/MS experiments were available at CPTAC public data portal (https://cptac-data-portal.georgetown.edu). MS/MS spectra were searched using the MSFragger database search tool [8] and further processed using the Philosopher pipeline [9] (<https://github.com/Nesvilab/philosopher>). The PSM files obtained from Philosopher pipeline were then processed using TMT-Integrator (<https://github.com/Nesvilab/TMT-Integrator>) to generate summary reports at the protein level.

Similar to the other studies, each TMT multiplex in the CPTAC CCRCC Discovery study contained a common internal reference sample, which was made through pooling equal amount of individual tumor samples in the study. This reference sample served as internal standards for all proteins and phosphoproteins quantified.

For each PSM of one TMT 10-plex, the intensity in each TMT channel was log2 transformed, and the reference channel intensity (pooled reference sample) was subtracted from the log2 intensity of the other nine channels (samples), thus generating relative intensity of each peptide at log2-based ratio-to-reference scale (referred to as ‘ratios’ below). PSMs were then grouped on the protein level after certain filtering, and the median was calculated from the remaining ratios to represent the ratio for each sample, for every protein. The Reference Intensity of one protein in one TMT-10plex was estimated using the sum of the MS1 intensities of the top three most intense peptide ions [10] in the TMT 10-plex. The overall Reference Intensity was then computed as the mean across all reference intensities in the study for each protein (with minimum imputation for the missing intensity values of reference channels). The final protein abundance of a sample was defined as the sum of intensity ratios and reference sample MS1 intensities at the protein level. In the end, for the control of sample-specific artificial effect, global normalization was applied to the protein abundance matrix by (1) centering the sample-wise median to the same level, and (2) scaling each sample with the sample-wise median absolute deviation (MAD). Batch effect of TMT 10-plexes was also removed by using Combat [11]. More detail on the MS1 intensity table generation for CCRCC study can be found in the study publication [4].

**A4. Decoyed Data Sets for Imputation Evaluation during the Challenge Phase.**

During the challenging phase, we created decoyed data sets with pseudo missing values based on the real proteomics data sets from CPTAC/TCGA Breast Cancer and CPTAC Prospective Ovarian Cancer studies [1,2] (**Table S1**). Specifically, training data sets were generated based on the global proteomics data from CPTAC/TCGA breast cancer study [1], while test sets were generated based on the global proteomics data from the CPTAC Prospective Ovarian Cancer Study [2] (**Table S1**). The latter were not publicly available during the Challenge Phase.

In each simulation run (training and testing), we started with the abundance matrix of the subset of proteins/genes with complete observation in all samples. In the following text, we refer to these matrices as Basis matrices. Specifically, the Basis matrix of the training data was from the CPTAC/TCGA Breast Cancer data of 80 tumor samples (77 patients). 7927 genes with complete observation were selected. The Basis matrix of the testing data was from the CPTAC Prospective Ovarian Cancer Data of 83 tumor samples of 80 patients (from the total 103 samples in the cohort). 8203 genes with complete observation were included.

Given the Basis matrix, in each simulation run, we randomly sampled a collection of data points according to probability models that mimic the missing mechanisms in labeled proteomics experiments and set the values of these data points to NA. In this way, we established the ground truth of each missing data point, and could use these ground truths for evaluation. Specifically, “biological” missing (true 0s) and “instrumental” missing were generated using different mechanisms. The biological missing events (1/0 variables) across different proteins were created to share the similar correlation structure as the protein abundances. We introduced 10% biological missing in the training data sets, while utilized a range of biological missing rates (1%, 2.5%, 5%, 10%) for the testing data sets to mimic the diverse range of real data scenarios. On the other hand, the instrumental missing events were created using the abundance dependent missing mechanisms [5] inferred from the CPTAC/TCGA breast cancer proteomics data set. Both instrumental missing and biological missing were annotated as ‘NA’ in the final training and testing data sets.

To facilitate the evaluation of the imputation algorithms, we generated in total 10 training data sets and 100 testing data sets for each of the leader board and final rounds during the Challenging phase (**Fig. 2a**).

**A5. Imputation Evaluation during the Community Phase**

During the community phase, to evaluate imputation performances, we utilized protein abundance profiles of two sets of tissue samples from the same group of tumors (n=32) in the CPTAC/TCGA ovarian study [3] (**Table S1**). Specifically, the two sets of samples were quantified separately by two proteomics labs: the Pacific Northwest National Laboratory (PNNL) and a proteomics lab from Johns Hopkins University (JHU). We referred to the resulting data sets as the PNNL-data (n=32) and the JHU-data (n=32) respectively. The normalized protein abundance tables for the PNNL- and JHU-data were derived as described in A.3. To further remove artificial “batch effects” associated with the two proteomics labs, the two abundance tables were aligned by shifting the mean abundance of each protein in the JHU data set to the same level as that in the PNNL data. Based on the final normalized and aligned abundance matrices, the average gene-wise Spearman correlation between the JHU and PNNL data is 0.95 (with SD of 0.039). Other quality assessment of the JHU and PNNL proteomics data sets were described in the original publication [3].

During the community phase, all imputation methods were firstly applied to the PNNL-data of 3027 genes (n=32). The imputed data points were then evaluated against the corresponding observed data points in JHU-data, which is regarded as a good approximation to the true values that were missing in the PNNL-data.

#### A6. Evaluation Metrics and Tie Breaking for Determining the Final Rank in the Challenge

The performance of imputation algorithms were evaluated based on Pearson Correlation coefficients (Cor) and the Normalized Root-Mean-Square Deviation (NRMSD) between imputed and the underlying true values of missing data points.[12] Specifically, denote *X=(x_i_)* as the imputed values, and *Y=(y_i_)* as the underlying true values. The Cor and NRMSD were calculated using the below formulas:

$Cor=\frac{1}{n_{missing}-1}\sum_{i=1}^{n_{missing}} \frac{(x_{i}-\bar{X})(y_{i}-\bar{Y})}{s_{x}s_{y}}$,

$NRMSD=\frac{\sqrt{\sum_{i=1}^{n_{missing}} {{(y_{i}-x_{i})}^{2}}/{n_{missing}}}}{y_{max}-y_{min}}$,

where $s_{x}=\left( \frac{1}{n_{missing}-1}\sum_{i=1}^{n_{missing}} (x_{i}-\bar{X})^{2} \right)^{1/2}$, and $s_{y}=\left( \frac{1}{n_{missing}-1}\sum_{i=1}^{n_{missing}} (y_{i}-\bar{Y})^{2} \right)^{1/2}.$ Note, for each protein, Cor was calculated based on instrumental missing spots, while NRMSD was calculated based on all missing spots.

Average Cor and NRMSD across the 100 testing data sets in the final round were used to identify the winning teams. The performance of the K-Nearest-Neighbor (KNN) imputation method was used as the reference. For three teams, *DMIS_PTG, BruinGo,* and *Jeremy*, their Cor scores were larger than that of KNN, while their NRMSD scores were smaller than or equal to that of KNN (**Fig 2b**). To better rank the performance of these three teams, we further compared the performances of each pair of teams using the below criteria:

1.*Confidence Intervals* For each team, we computed 95% Confidence Intervals (CI) of the Cor and NRMSD scores based on the results from the 100 test data sets. We declared two teams’ performances were statistically different, if one team had all CIs higher than and not overlapping with that of the other team. Since different biological missing rates shall lead to different levels of evaluation scores, to make the CI estimates more meaningful, we calculated CIs of Cor and NRMSD for the four groups with different biological missing rates separately. We then counted for how many of the 4 scenarios one team appeared to be better than another team.

2. *Bayes Factor* For a pair of teams, we estimated their Bayes Factor (BF) [6,12] using their performance across 100 test data sets. We first counted the number of test data sets on which team A performed better than team B. Denote this number as a. We then counted the number of test data sets on which team A performed worse than team B. Then the BF was calculated as a/(a+b). Two teams were declared to perform differently if the Bayes Factor is larger than 10 or smaller than 0.1.

CIs based comparisons using either Cor or NRMSD were summarized in **Table S2**. A number of “4” suggests that the scores of the row team were significantly better than that of the column team across all 4 levels of biological missing rate settings. The Cor based evaluation (**Table S2a**) suggested an order of *DMIS_PTG>Jeremy>BruinGo.* The NRMSD based evaluation, however, was not very informative (**Table S2b**),

BF based comparison was summarized in **Table S3**. A number larger than 10 in the table suggests that the performance of the row team is significantly better than that of the column team. If the number is smaller than 0.1, then the performance of the row team was viewed to be significantly worse than that of the column team. The Cor based evaluation (**Table S3a**), again, suggested an order of *DMIS_PTG>Jeremy>BruinGo.* The NRMSD based evaluation suggested *DMIS_PTG* is better than the other teams. In conclusion, team DMIS_PTG won this sub-challenge*.*

**Table S2. Comparing Results Using Confidence Intervals.** Each value in the table represents the number of settings in which the CI of the row team was strictly higher than that of the column team. (a) is showing comparison based on Cor, and (b) is showing comparison based on NRMSD.

**(a).**

| **Correlation Confidence Interval** | **R1** | **R2** | **R3** | **Ref** |
| --- | --- | --- | --- | --- |
| DMIS_PTG(R1) |  | 4 | 4 | 4 |
| BruinGo(R2) | 0 |  | 0 | 0 |
| Jeremy(R3) | 0 | 4 |  | 4 |
| KNN(Ref) | 0 | 0 | 0 |  |

**(b).**

| **NRMSD Confidence Interval** | **R1** | **R2** | **R3** | **Ref** |
| --- | --- | --- | --- | --- |
| DMIS_PTG(R1) |  | 0 | 0 | 0 |
| BruinGo(R2) | 0 |  | 0 | 0 |
| Jeremy(R3) | 0 | 0 |  | 0 |
| KNN(Ref) | 0 | 0 | 0 |  |

**Table S3. Comparing results Using Bayes Factors.** Each value in the table represents the Bayes Factor of the row team vs. the column team based on either Cor (a) or NRMSD (b).

**(a)**

| **Correlation Bayes Factor** | **R1** | **R2** | **R3** | **Ref** |
| --- | --- | --- | --- | --- |
| DMIS_PTG(R1) |  | Inf | Inf | Inf |
| BruinGo(R2) | 0 |  | 0 | 1.04 |
| Jeremy(R3) | 0 | Inf |  | Inf |
| KNN(Ref) | 0 | 0.96 | 0 |  |

**(b)**

| **NRMSD BayesFactor** | **R1** | **R2** | **R3** | **Ref** |
| --- | --- | --- | --- | --- |
| DMIS_PTG(R1) |  | Inf | Inf | Inf |
| BruinGo(R2) | 0 |  | 1 | 1.38 |
| Jeremy(R3) | 0 | 1 |  | 1 |
| KNN(Ref) | 0 | 0.72 | 1 |  |

#### A7. Protein Clusters for Imputation Evaluation in the Community Phase

To understand how imputation performance may be affected by protein characteristics, we summarized the imputation performances for stratified protein clusters. We utilized three different criteria to define protein clusters: **protein closeness**, **pseudo missing performance**, and **protein abundance**. Detailed definition of protein clusters are provided in below:

1. **Protein Closeness:** Consider the subset of proteins having at least one missing data point in the PNNL data. For a given protein, we first identify 50 other proteins whose abundances showed the largest correlation with that of the given protein. We then calculated the closeness score of the given protein as the average correlation between each of the 50 selected proteins and the given protein. Based on the closeness scores, we split 289 proteins that were eligible for evaluation into 4 clusters of roughly equal sizes. The clusters were ordered according to their closeness scores, such that the 1^st^ cluster contains proteins with the lowest closeness scores, while the 4^th^ cluster contains the ones with the highest closeness scores.
2. **Pseudo missing performance:** During the bagging step, pseudo missing events were introduced to the real data set (the PNNL-data) to generate a collection of perturbed data sets. Since the underlying truths (observed data points in the original PNNL-data) were established by these pseudo missing data points, their imputed values in the output of the bagging step were used to evaluate the imputation performance. Specifically, for each protein, pseudo-missing NRMSD was calculated by comparing the imputed values of the pseudo missing data points of this protein across all the bagging data sets and the corresponding observed data points in the original PNNL-data. We then used these pseudo-missing NRMSDs to divide the 289 proteins into four clusters. These clusters were ordered from low performance to high performance, meaning that the 1^st^ cluster had the highest pseudo-missing NRMSD (the lowest performance); and the 4^th^ cluster has the lowest pseudo-missing NRMSD (the highest performance).
3. **Protein abundance:** Finally, we defined protein clusters based on observed mean protein abundances. Specifically, the 1^st^ cluster contains proteins with the lowest mean abundances, while the 4^th^ cluster contains proteins with the highest mean abundances.

#### A8. Baseline Algorithms used in the DreamAI.

#### The consensus imputation strategy of DreamAI includes three baselines algorithms: ADMIN, knn.impute, and missForest. The details of each baseline imputation algorithm are provided in below.

***ADMIN***

This method was designed for imputation of isotopic labeling proteomics data in which batch effects exist and missing data is dependent on protein abundances [13]. In this manuscript, we refer to this method as ADMIN (Abundance dependent Missing Data Imputation). For a given protein, ADMIN models its abundance as a linear combination of the abundances of the closest K neighbors of the protein. Specifically, a mixed effects model is used and batch effects in the data are characterized through a random effect in the model. The closest neighbors are determined by the pairwise Pearson correlation. At the same time, a probability model is utilized to characterize the abundance dependent missing mechanism: missing rate is modeled to be exponentially linearly correlated with the ‘true’ abundances. ADMIN then utilizes an EM(expectation- maximization) algorithm to estimate the linear prediction model and the abundance dependent missing model. The expectation of the missing data points based on the estimated model are returned as the final imputation result. To avoid huge computation consumption, the default number of neighbors in algorithm is set to 10.

***knn.impute***

knn.impute is a function designed to impute missing values in gene expression data sets based on K-nearest neighbor averaging [14,15]. For each gene with missing values, K nearest neighbors were found using a Euclidean distance metric, confined to the columns for which that gene is NOT missing. After the k nearest neighbors are identified, imputed value of a missing element is the average of those (non-missing) elements of its neighbors. To increase computation efficiency, gene sets over a certain threshold (set as 1500 in the package) were broken into blocks using two-mean clustering. This is done recursively till all blocks have less than the max number of genes. For each block, k-nearest neighbor imputation is done separately.

**missForest**

missForest is developed to impute missing values particularly in mixed-type data: continuous and/or categorical data including complex interactions and nonlinear relations.[16] missForest uses an iterative imputation scheme: (1) train a Random forest model on observed values; (2) predict the missing values; and (3) repeat (1) and (2) until the model converges. Random forest is chosen to model the missing value as it can handle mixed-type data and is known to perform very well under conditions like high dimensions, complex interactions and non-linear data structures. In case of high-dimensional data some parameters in the algorithm are suggested with a relatively small value, for example: number of trees to grow in each forest and number of variables randomly sampled at each split to obtain an appropriate imputation result within a feasible amount of time. Moreover, it can be run parallel to save computation time using an appropriate backend.

#### A9. Methods of the Top Three Teams

**SpectroFM: Matrix factorization-based imputation**

In the computer science domain, the imputation of missing values, which has been the focus of many studies, can be considered as a recommendation task since a user’s unobserved preferences are represented as missing values in a user-item matrix. Given a user-item matrix, a recommendation system predicts a user’s preferences for an item based on other users’ existing preferences for the item and the user’s preferences for other items. This is analogous to the task in this challenge. If we consider proteins as items and patients as users, it is possible to exploit collaborative filtering algorithms. We first apply Z-normalization to a protein abundance data matrix to make the data fit a normal distribution. We save the mean and variance to revert the data to its original scale when we perform imputation. We train a low-rank matrix factorization model on existing values in the normalized abundance matrix. For the implementation of the matrix factorization model, we use LibFM, a factorization machine library [17]. Using the calculated latent parameter matrix of proteins and the latent parameter matrix of patients in the model, we reconstruct the best approximation of the original input matrix by multiplying the two latent matrices. Since the latent matrices are dense, the missing values in the original matrix are imputed in the reconstructed approximated matrix. We set the dimensions of the latent protein and patient matrices to 40. Consequently, the rank of the reconstructed approximated matrix is 40. We use a Markov chain Monte Carlo (MCMC)[18] algorithm to optimize parameters. One of the advantages of MCMC is that it integrates regularization parameters into the model, which allows us to skip hyper parameter optimization. After the imputation of missing values by the multiplication of the latent matrices, we revert the normalized values to their original scale using the saved mean and variance. More details of the algorithm were provided int the below pseudo code.

**Algorithm 1**: SpectroFM

**Input**: Binary missing indicator feature vector x and observed protein abundance values y^obs^

**Output**: imputed protein abundance values y^miss^

Initialize model parameter θ

Z-Normalize y values to ỹ using

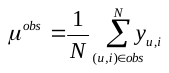

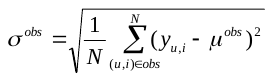

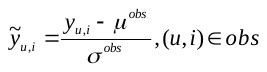

Obtaini optimal parameter:

For t in 1,...,7

1: θ^*^ = MCMCOptimizer(x, ỹ, θ)

2: θ = θ^*^

Impute the missing values ỹ_u,i_^miss^ using

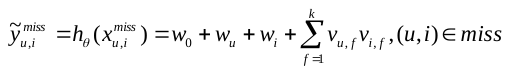

Return ỹ^miss^ to original scale values y^miss^ using

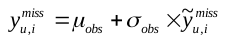

**RegImpute: Regression-based imputation**

A conventional, post-processed proteomics dataset usually takes the form of a two-dimensional array. From the perspective of training a regression model, the columns of an array can be interpreted as features (dimensions), and the rows can be considered as training instances (or vice-versa). The features and instances can be used to train a predictive model to impute unobserved contains missing values. One solution is to divide data sets into subsets, on which models can be trained. However, this approach can be very time consuming. A second approach is to train a model on only complete dataset without missing values. The drawback of this approach is that samples with missing values may be characteristically different from samples without missing values (e.g., not missing at random (NMAR) versus missing at random (MAR)). RegImpute is a combination of the two approaches above and uses a simple imputation method such as mean imputation on the existing values to generate a complete training set. In addition, users can impute missing values using the values (e.g., zeros) selected by the users. Then, we use ridge regression, which is a fast and robust linear regression technique. Ridge regression is an extension of linear regression, and its regularization prevents it from overfitting. Ridge regression performs regularization by adjusting weights to avoid focusing on only a few features [19]. Using single regression on the dataset may be sufficient if the initial guesses are nearly correct or if there are few missing values. However, in some cases, the initial regression values are heavily influenced by a prior assumption(s). For this reason, performing regression several times may reduce estimation errors. At each iteration, we use the imputed missing values from the previous imputation to improve regression for the current imputation. At some point, usually after ~10 iterations, convergence is reached.

**Algorithm 2**: RegImpute

**Input:** *data matrix Y of protein abundance*

**Output:** Y^*^

For each iteration n:

1. For each column Y*_i_* in data Y (i = 1,...,n) Split the data into two subsets:

Y_miss,i_: rows with Y*_i_* missing

Y_obs,i_: rows with Y*_i_* observed

2 Fill NAs in Y_obs,i_ with the imputed values of Y_obs,i_ in iteration n-1 (fill in 0s if n=1)

3 Train ridge regression model on data Y_obs,i_, to associate ith column with all the other columns, obtain β to solve the minimization:

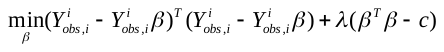

4 Fill in NAs in Y_miss,i_ with the imputed values of Y_obs,i_ in iteration n-1 (fill in 0s if n=1)

for all but not ith column

5 Use trained model on Y_obs,i_ to predict ith column in Y_miss,i_ with the other columns as predictor

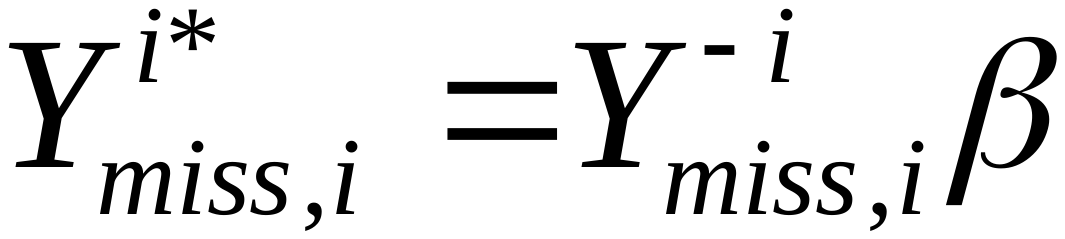

6 repeat 1 to 5 until convergence of imputed value.

**Birnn: Matrix completion and Bagging-based imputation**

We consider the imputation of missing protein abundances in a protein-sample matrix as a matrix completion problem. We assume that all the protein abundances have the same data distribution because they are from the same type of cancer, and thus the matrix is assumed to have a low rank structure. Based on this assumption, we used the iteratively reweighted nuclear norm (IRNN)[20] algorithm with the smoothly clipped absolute deviation (SCAD)[21] penalty, which is a non-convex penalty function on singular values, to better approximate the rank function and enhance low rank matrix approximation. Moreover, we use the bootstrap aggregating algorithm to train multiple models on sampled sub-datasets of the original dataset. The final prediction is given by aggregating the outputs of the multiple models. The bootstrap aggregating algorithm can help prevent models from over fitting by reducing model variance, which contributes to performance improvement.

**Algorithm 3**: Birnn

**Initialize:** *k* = 0, X^k^, w^k^_i_, i = 1,2,...,min(m,n)

**Output:** X^*^

1. while not converge do
2. Update X^k^ by solving

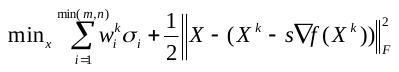

with **Weighted Singular Value Thresholding (WSVT).**

Update the weights w^k^_i_, i = 1,2,...,min(m,n) by

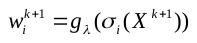

where

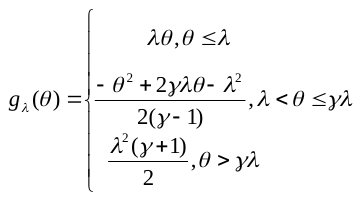

3. end while

#### A10. The NCI-CPTAC Proteogenomics DREAM imputation Challenge Participants

#### Table S4. List of NCI-CPTAC Proteogenomics DREAM imputation Challenge Participants.

| **Full Name** | **Affiliation** |
| --- | --- |
| Gajendra Pal Singh Raghava | Indraprastha Institute of Information Technology, Delhi, INDIA |
| Sunil V Kalmady | University of Alberta,Edmonton, Alberta, Canada |
| Harpreet Kaur | CSIR-Institute of Microbial Technology, Chandigarh, INDIA |
| Piyush Agrawal | CSIR-Institute of Microbial Technology, Chandigarh, INDIA |
| Salman Sadullah Usmani | CSIR-Institute of Microbial Technology, Chandigarh, INDIA |
| Eunji Heo | Deargen Inc. & School of Computing, KAIST, Daejeon, South Korea. |
| Bora Lee | Deargen Inc., Daejeon, South Korea |
| Yunpeng Liu | Department of Biology, Massachusetts Institute of Technology, Cambridge MA, USA |
| Wei Chen, | Department of Biology, Southern University of Science and Technology, Shenzhen, China. |
| Yue Shan | Department of Biostatistics, The University of North Carolina at Chapel Hill, USA |
| Hongtu Zhu | Department of Biostatistics, the University of Texas MD Anderson Cancer Center, USA |
| Kaixian Yu | Department of Biostatistics, the University of Texas MD Anderson Cancer Center, USA |
| Hongyang Li | Department of Computational Medicine and Bioinformatics, University of Michigan, Ann Arbor, MI, USA |
| Yuanfang Guan | Department of Computational Medicine and Bioinformatics, University of Michigan, Ann Arbor, MI, USA |
| Jaewoo Kang | Department of Computer Science and Engineering & Interdisciplinary Graduate Program in Bioinformatics, College of Informatics, Korea University |
| Daehan Kim | Department of Computer Science and Engineering, College of Informatics, Korea University |
| Keonwoo Kim | Department of Computer Science and Engineering, College of Informatics, Korea University |
| Minji Jeon | Department of Computer Science and Engineering, College of Informatics, Korea University |
| Sunkyu Kim | Department of Computer Science and Engineering, College of Informatics, Korea University |
| Yonghwa Choi | Department of Computer Science and Engineering, College of Informatics, Korea University |
| Tengfei Li | Department of Radiology, The University of North Carolina at Chapel Hill, USA |
| Liuqing Yang | Department of Statistics and Operations Research, The University of North Carolina at Chapel Hill, USA |
| Maomao Ding | Department of Statistics, Rice University, USA |
| Jingyi Jessica Li | Department of Statistics, University of California, Los Angeles, CA, USA |
| Kexin Li | Department of Statistics, University of California, Los Angeles, CA, USA |
| Xinzhou Ge | Department of Statistics, University of California, Los Angeles, CA, USA |
| Huiyuan Chen, | Electrical Engineering and Computer Science, Case Western Reserve University, USA |
| Ke Hu, | Electrical Engineering and Computer Science, Case Western Reserve University, USA |
| Kumardeep Chaudhary | Epidemiology Program, University of Hawaii Cancer Center, Honolulu, HI, 96813, USA |
| Nai-Wen Chang | Graduate Institute of Biomedical Electronics and Bioinformatics, National Taiwan University, Taipei, Taiwan |
| Jia Xin Yu, | Icahn School of Medicine at Mount Sinai, New York, New York |
| Devishi Kesar | Indraprastha Institute of Information Technology, Delhi, INDIA |
| Sherry Bhalla | Indraprastha Institute of Information Technology, Delhi, INDIA |
| Mehreen Ali | Institute for Molecular Medicine Finland, Helsinki Institute of Life Science, University of Helsinki, Finland |
| Ábel Fóthi | Institute of Enzymology, Research Centre for Natural Sciences, Hungarian Academy of Sciences, Budapest, Hungary |
| Ching-Tai Chen | Institute of Information Science, Academia Sinica, Taipei, Taiwan |
| Ting-Yi Sung | Institute of Information Science, Academia Sinica, Taipei, Taiwan |
| Heewon Lee | Interdisciplinary Graduate Program in Bioinformatics, College of Informatics, Korea University |
| Hwisang Jeon | Interdisciplinary Graduate Program in Bioinformatics, College of Informatics, Korea University |
| Sandeep Kumar Dhanda | La Jolla Institute for Immunology, La Jolla, CA, USA |
| Swapnil Mahajan | La Jolla Institute for Immunology, La Jolla, CA, USA |
| San-Yuan Wang | Master Program in Clinical Pharmacogenomics and Pharmacoproteomics, College of Pharmacy, Taipei Medical University, Taipei, Taiwan |
| Shujiro Okuda | Niigata University, Niigata, Japan |
| Yasuhiro Kambara | Niigata University, Niigata, Japan |
| Laura L. Elo | Turku Bioscience Centre, University of Turku and Åbo Akademi University, Turku, Finland |
| Mehrad Mahmoudian | Turku Bioscience Centre, University of Turku and Åbo Akademi University, Turku, Finland |
| Sohrab Saraei | Turku Bioscience Centre, University of Turku and Åbo Akademi University, Turku, Finland |
| Tomi Suomi | Turku Bioscience Centre, University of Turku and Åbo Akademi University, Turku, Finland |
| Tommi Välikangas | Turku Bioscience Centre, University of Turku and Åbo Akademi University, Turku, Finland |
| Russell Greiner | University of Alberta, Edmonton, Alberta, Canada |
| Roberto Vega | University of Alberta,Edmonton, Alberta, Canada |
| Jeremy R. Jacobsen | University of Colorado Boulder, Boulder, Colorado, USA |

7. CPTAC Common Data Analysis Pipeline, <https://proteomics.cancer.gov/data-portal/about/common-data-analysis-pipeline>

8. Kong, Andy T., Felipe V. Leprevost, Dmitry M. Avtonomov, Dattatreya Mellacheruvu, and Alexey I. Nesvizhskii. "MSFragger: ultrafast and comprehensive peptide identification in mass spectrometry–based proteomics." *Nature methods* 14, no. 5 (2017): 513-520.
9. da Veiga Leprevost, Felipe, Sarah E. Haynes, Dmitry M. Avtonomov, Hui-Yin Chang, Avinash K. Shanmugam, Dattatreya Mellacheruvu, Andy T. Kong, and Alexey I. Nesvizhskii. "Philosopher: a versatile toolkit for shotgun proteomics data analysis." *Nature methods* 17, no. 9 (2020): 869-870.
10. Ning, Kang, Damian Fermin, and Alexey I. Nesvizhskii. "Comparative analysis of different label-free mass spectrometry based protein abundance estimates and their correlation with RNA-Seq gene expression data." *Journal of proteome research* 11, no. 4 (2012): 2261-2271.
11. Johnson, W. Evan, Cheng Li, and Ariel Rabinovic. "Adjusting batch effects in microarray expression data using empirical Bayes methods." *Biostatistics* 8, no. 1 (2007): 118-127.
12. Menden, Michael P., Dennis Wang, Mike J. Mason, Bence Szalai, Krishna C. Bulusu, Yuanfang Guan, Thomas Yu et al. "Community assessment to advance computational prediction of cancer drug combinations in a pharmacogenomic screen." *Nature communications* 10, no. 1 (2019): 1-17.
13. Wang, Minghui, Noam D. Beckmann, Panos Roussos, Erming Wang, Xianxiao Zhou, Qian Wang, Chen Ming et al. "The Mount Sinai cohort of large-scale genomic, transcriptomic and proteomic data in Alzheimer's disease." *Scientific data* 5 (2018): 180185.
14. Hastie, Trevor, Robert Tibshirani, Gavin Sherlock, Michael Eisen, Patrick Brown, and David Botstein. "Imputing missing data for gene expression arrays." (1999).
15. Troyanskaya, Olga, Michael Cantor, Gavin Sherlock, Pat Brown, Trevor Hastie, Robert Tibshirani, David Botstein, and Russ B. Altman. "Missing value estimation methods for DNA microarrays." *Bioinformatics* 17, no. 6 (2001): 520-525.
16. Stekhoven, Daniel J., and Peter Bühlmann. "MissForest—non-parametric missing value imputation for mixed-type data." *Bioinformatics* 28, no. 1 (2011): 112-118.
17. Rendle, Steffen. "Factorization machines with libfm." *ACM Transactions on Intelligent Systems and Technology (TIST)* 3, no. 3 (2012): 57.
18. Salakhutdinov, Ruslan, and Andriy Mnih. "Bayesian probabilistic matrix factorization using Markov chain Monte Carlo." In *Proceedings of the 25th international conference on Machine learning*, pp. 880-887. ACM, 2008.
19. Tikhonov, Andrei Nikolaevich, A. V. Goncharsky, V. V. Stepanov, and Anatoly G. Yagola. *Numerical methods for the solution of ill-posed problems*. Vol. 328. Springer Science & Business Media, 2013.
20. Lu, Canyi, Jinhui Tang, Shuicheng Yan, and Zhouchen Lin. "Generalized nonconvex nonsmooth low-rank minimization." In *Proceedings of the IEEE conference on computer vision and pattern recognition*, pp. 4130-4137. 2014.
21. Fan, Jianqing, and Runze Li. "Variable selection via nonconcave penalized likelihood and its oracle properties." *Journal of the American statistical Association* 96, no. 456 (2001): 1348-1360.

### Supplement B: SUPPLEMENTAL FIGURES

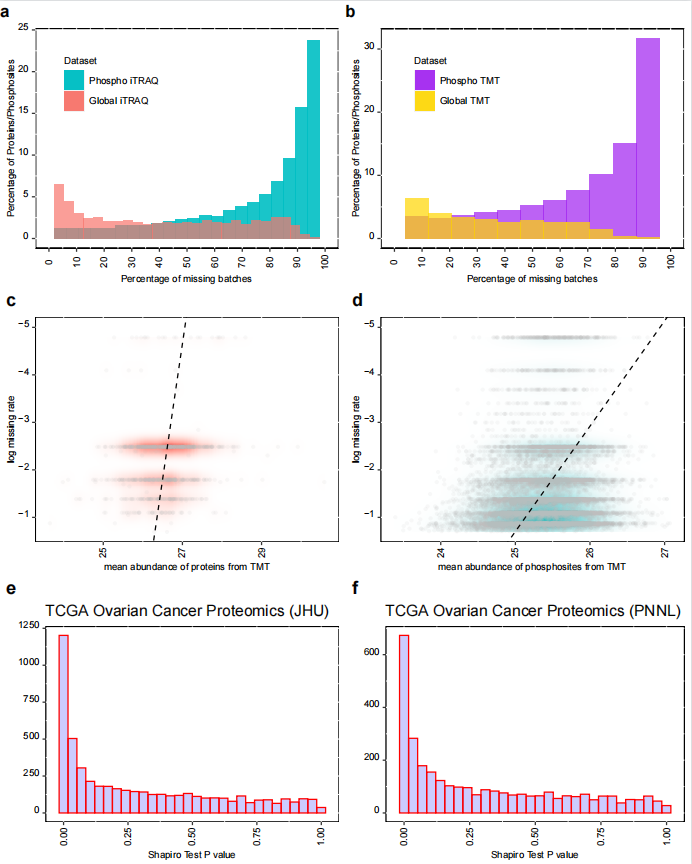

**Figure S1. Missing patterns and normality assessment of the global- and phosphor- proteomics data from CPTAC/TCGA Ovarian Cancer Study and the CPTAC Prospective Ovarian Cancer Study[2,3].** **(a-b)** Distributions of protein- and phosphosite-level (multiplex-)missing rates in global- and phospho-proteomics data sets from iTRAQ (a) and TMT (b) experiments respectively. **(c-d)** Scatter plots of protein-level or phosphosite-level missing rates vs. mean protein or phosphosite abundances based on observed data in the TMT global- or phospho- proteomics data sets. **(e-f)** Distributions of p-values from normality tests (Shapiro) on individual protein abundance distributions in the JHU-data or PNNL-data from the CPTAC/TCGA Ovarian Cancer Study.

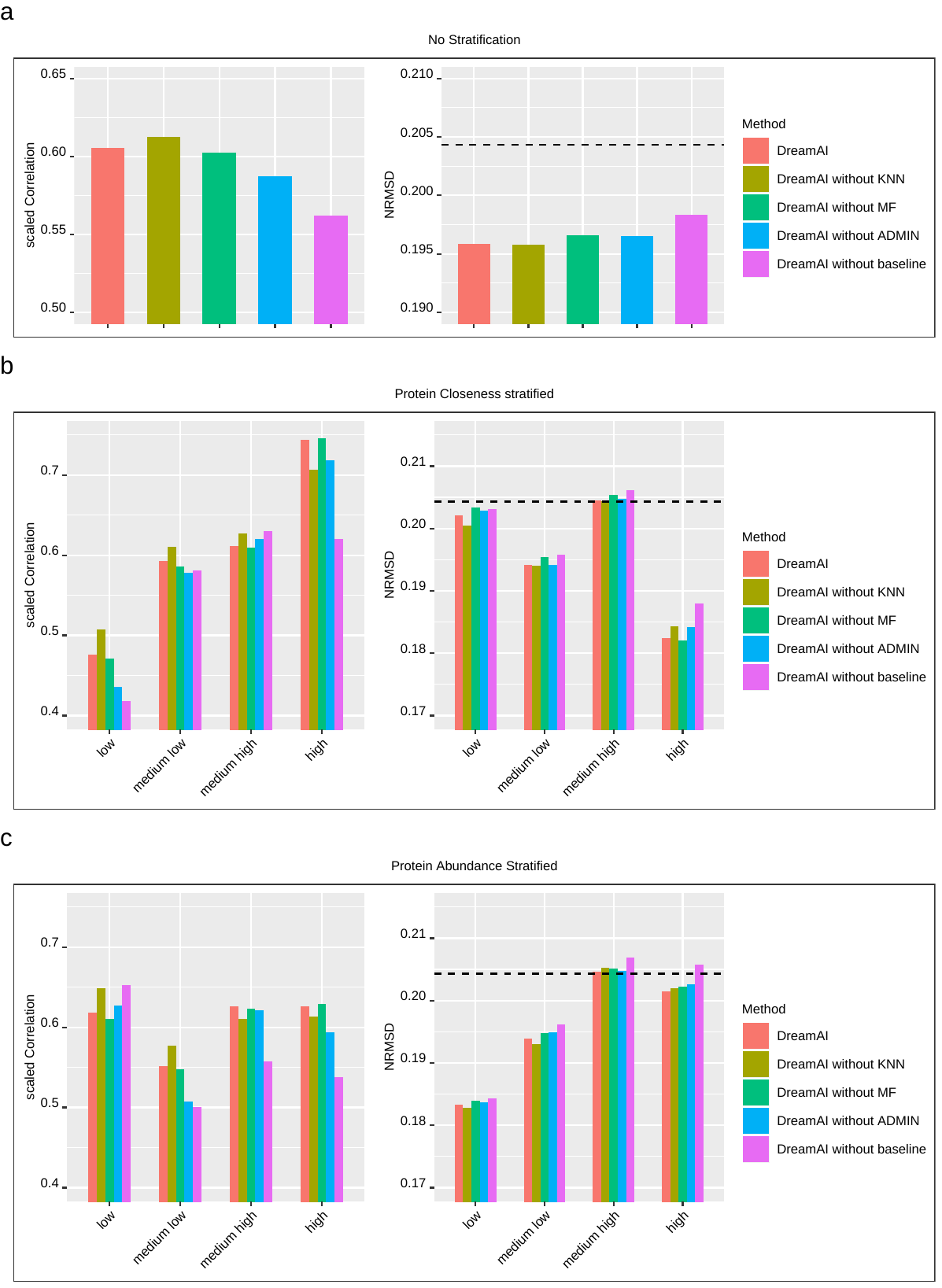

**Figure S2.** **Imputation performance of DreamAI with absence of one or all baseline methods** **on CPTAC/TCGA ovarian cancer data set.** **(a)** Average imputation performance (scaled Correlation and NRMSD) of all proteins. **(b-c)** Average imputation performance of different protein groups stratified by protein closeness (b) or protein abundances (c).

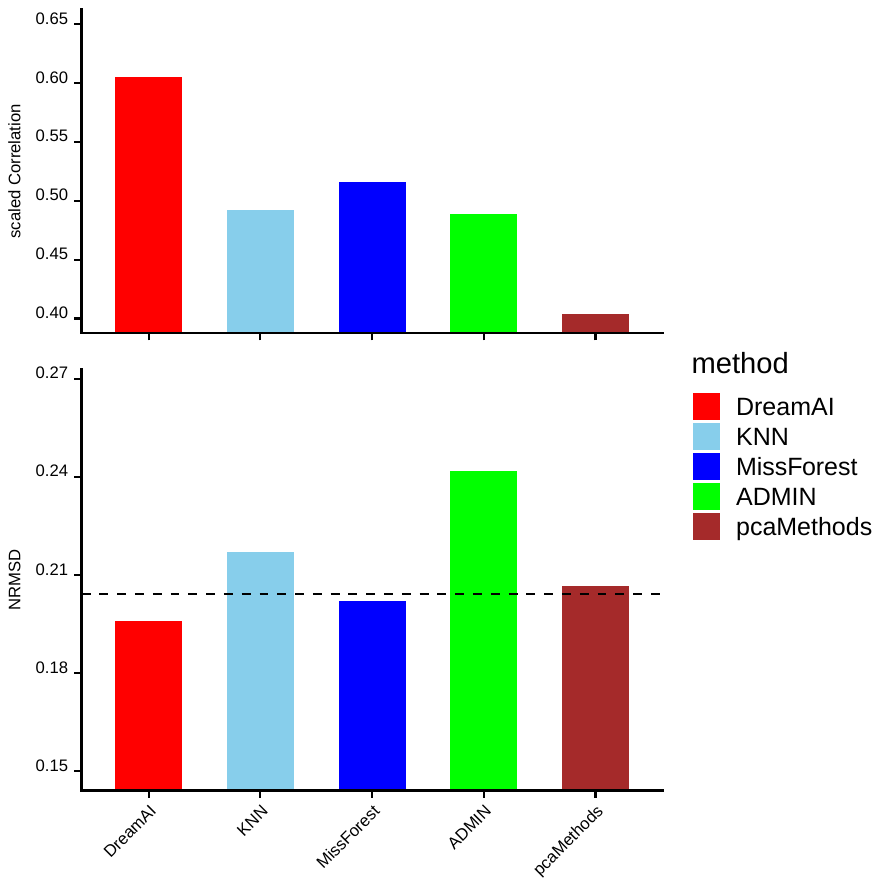

**Figure S3. Imputation performance of baseline methods on CPTAC/TCGA ovarian cancer data set**. Scaled Correlation was compared on the top panel and NRMSD was compared on the bottom panel. The dashed line in the bottom panel represents the background level of NRMSD between PNNL- and JHU-data based on data points observed in both data sets.

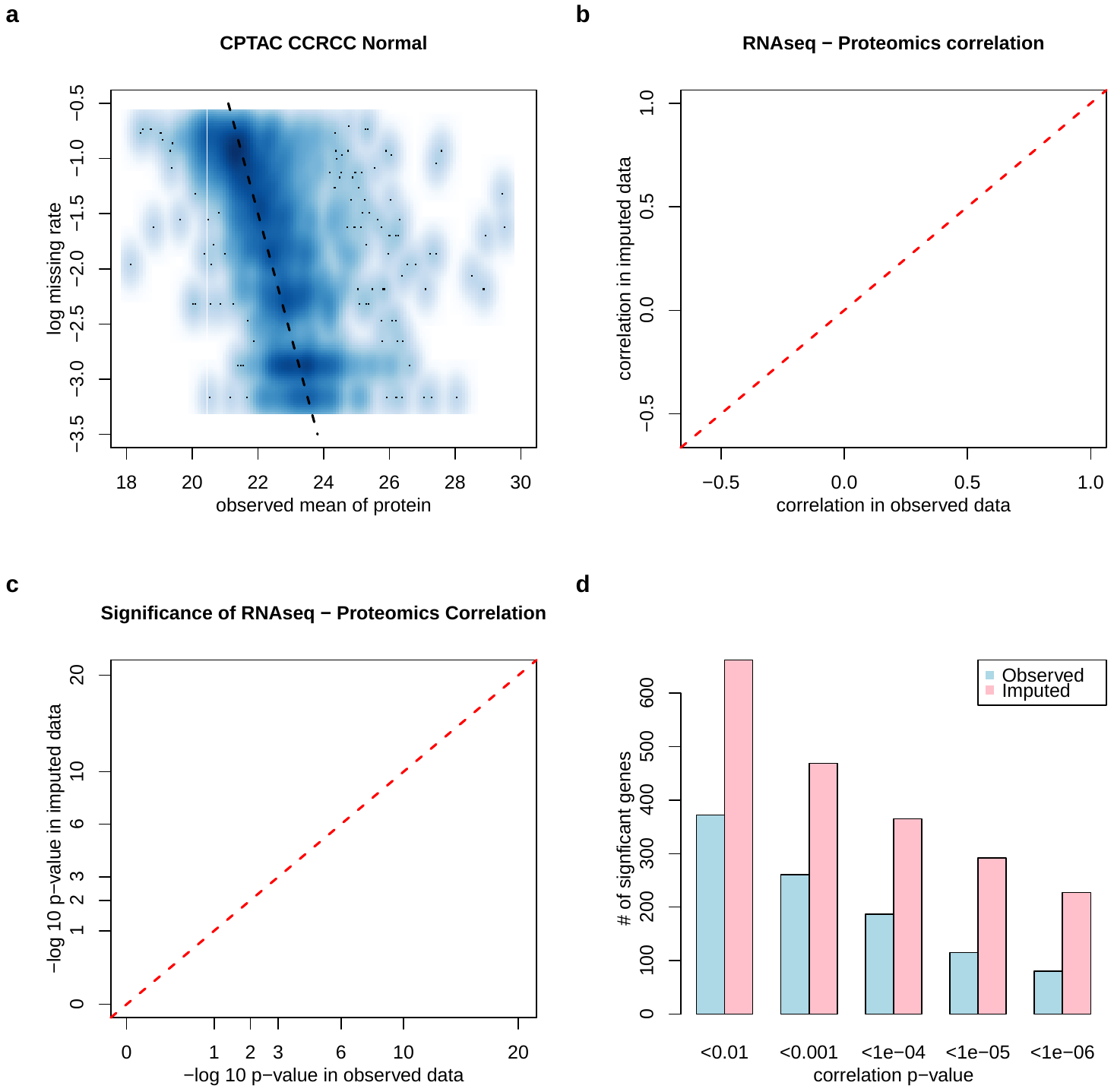

**Figure S4.** **For the CPTAC CCRCC NAT samples, proteomics data with** **DreamAI imputation shows improved concordance with their corresponding transcriptomics data.** **(a)** Scatter plot of protein-level missing rates vs. mean protein abundances based on observed values in the global proteomics data of 80 CCRCC NAT samples [4]. **(b)** Scatter plot of protein-RNA Spearman correlation based on the proteomics data with imputation (y-axis) vs. that without imputation (x-axis). **(c)** Scatter plot of significance levels (- log_10_p-value) of correlation test for protein-RNA association based on proteomics data with imputation (y-axis) vs. that without imputation (x-axis). **(d)** Number of genes showing significant protein-RNA association based on proteomics data with imputation (pink) or without imputation (blue) at different significance cutoff levels.

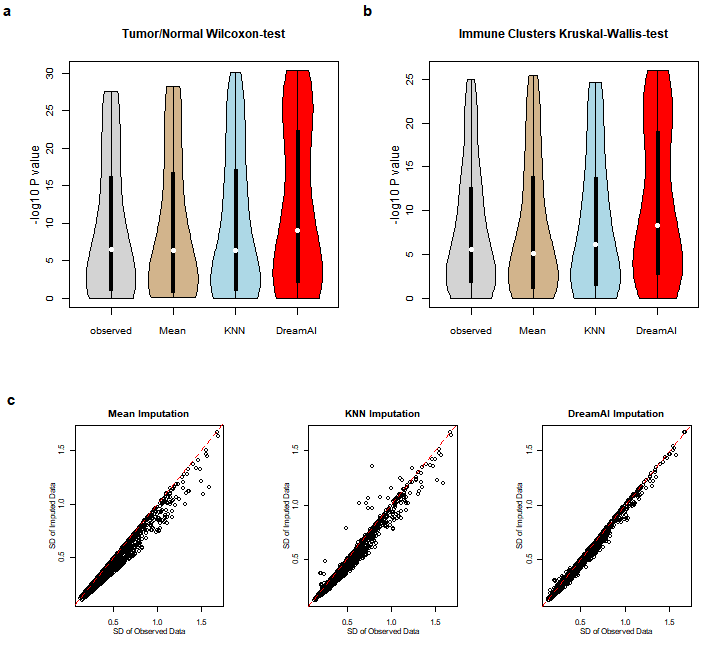

**Figure S5.** **Improved power to detect proteins associated with tumor/normal status or immune subtypes on the CPTAC CCRCC proteomic data with imputation by DreamAI. (a-b)** Distribution of p-values from association tests based on 4 different proteomics data matrices: without imputation (grey), mean value imputed (tan), KNN imputed (light blue) and DreamAI imputed (red). Two-sample Wilcoxon-tests were used for assessing association with tumor and NAT status (a) and Kruskal-Wallis tests were used for association with immune subtypes (b). Note, these plots focus on a subset of 92 proteins with substantially different imputed values between DreamAI and KNN (NRMSD>0.2). **(c)** Scatter plots comparing standard deviations of protein abundances before and after imputation. The three panels are for three different imputation strategies: Mean imputation (left), KNN imputation (middle) and DreamAI (right).
