## Supplementary figures and images for "DreamAI: algorithm for the imputation of proteomics data"

### Supp Fig 1

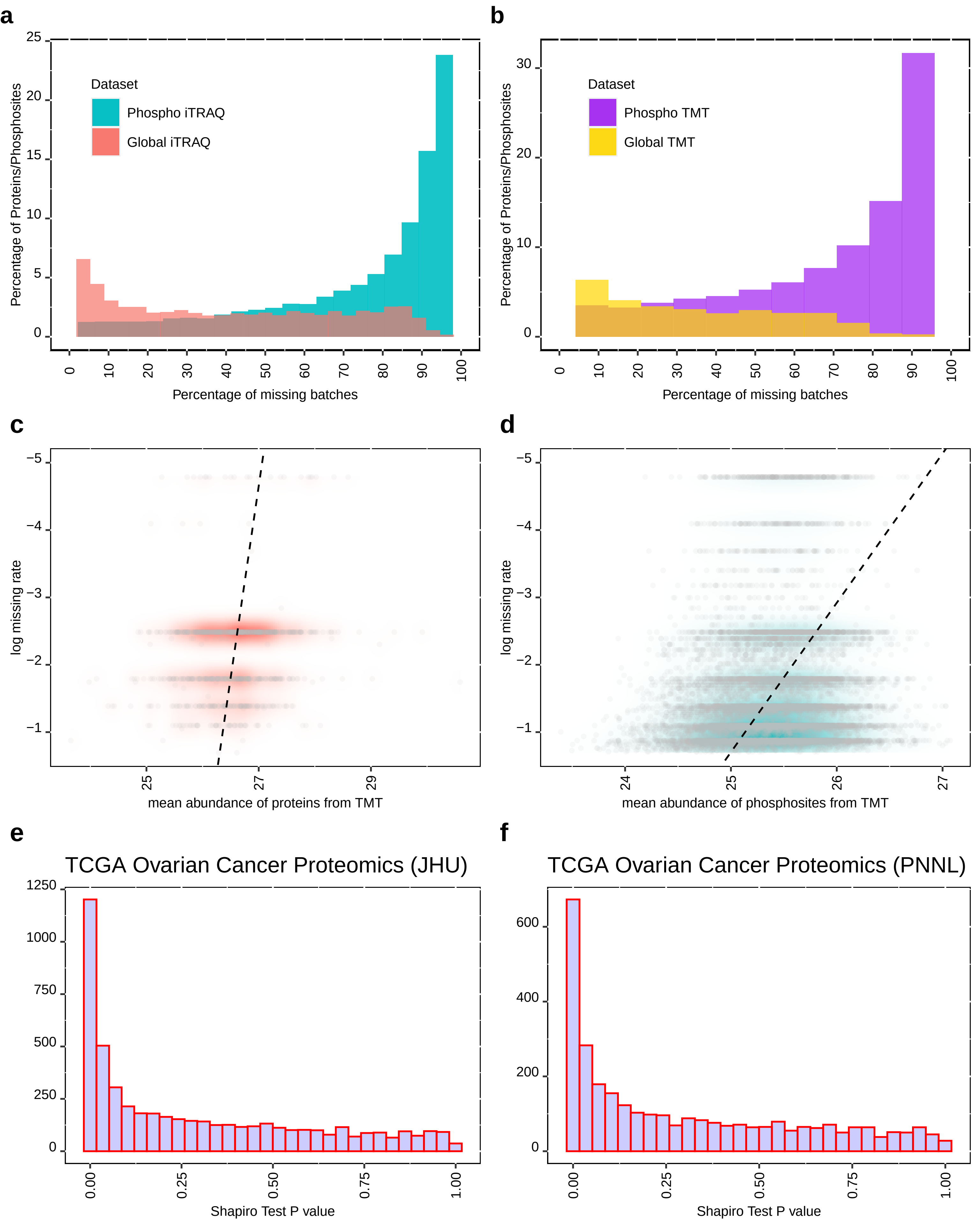

### Supp Fig 2

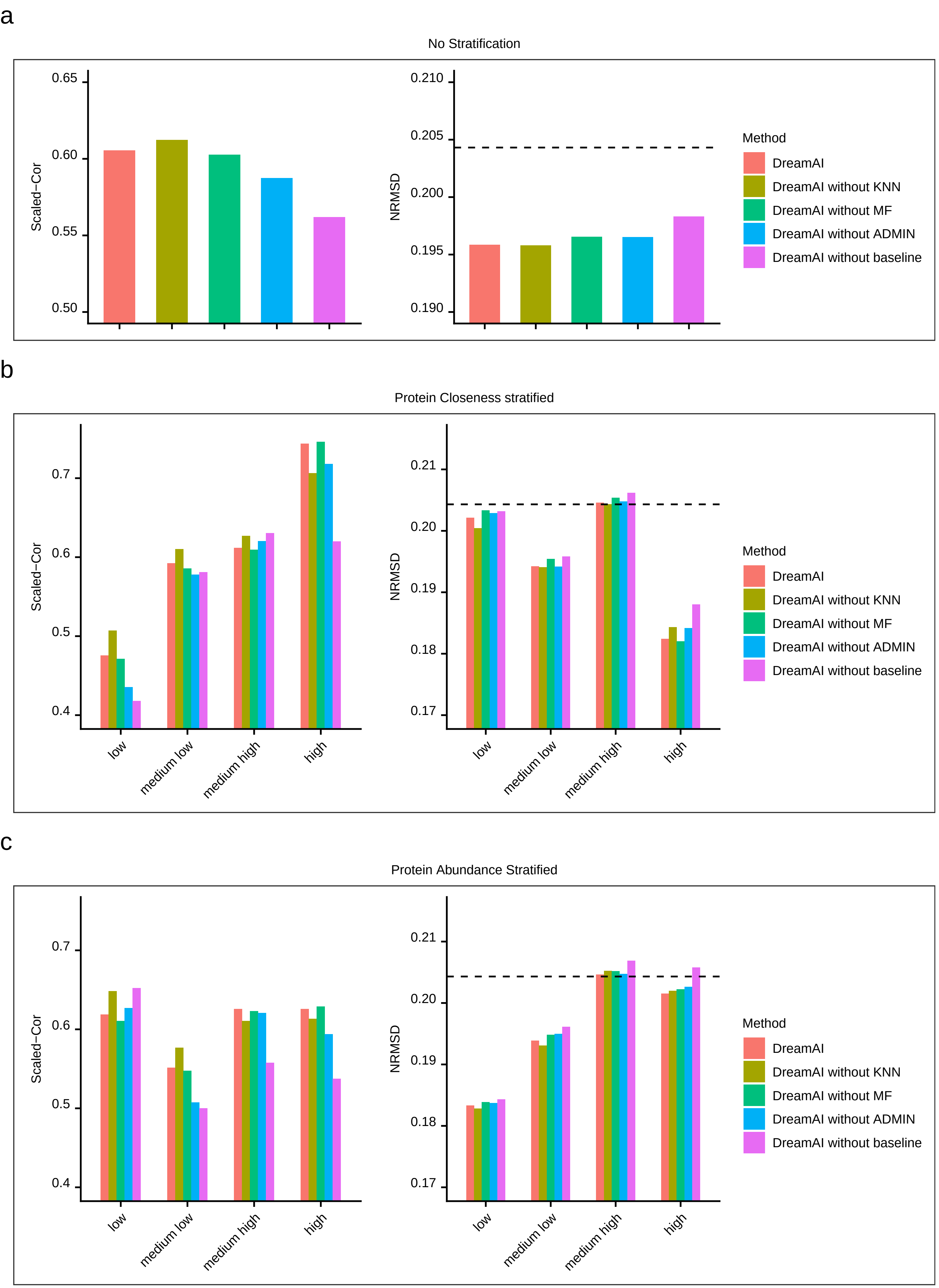

### Supp Fig 3

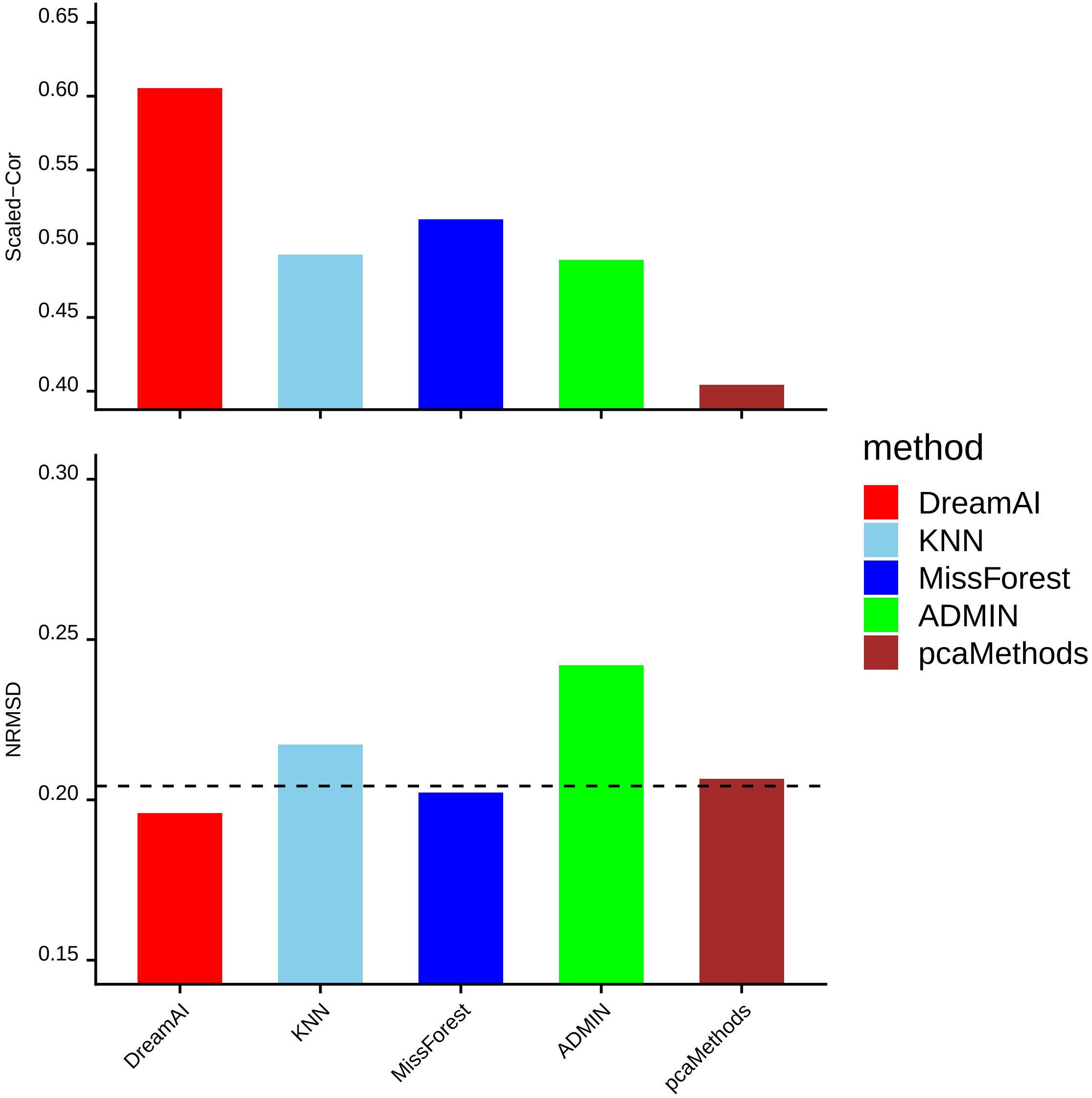

### Supp Fig 4

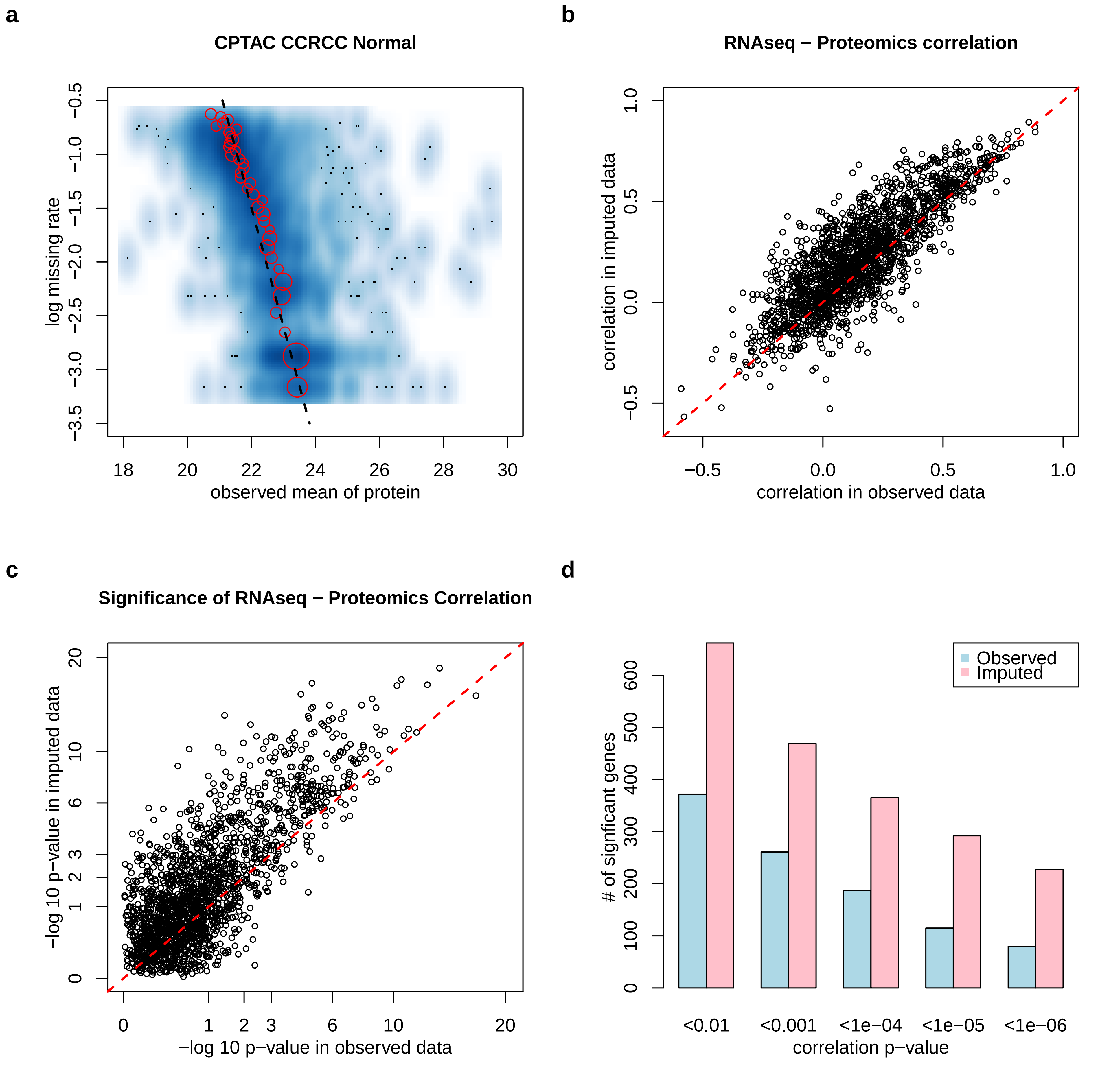

### Supp Fig 5

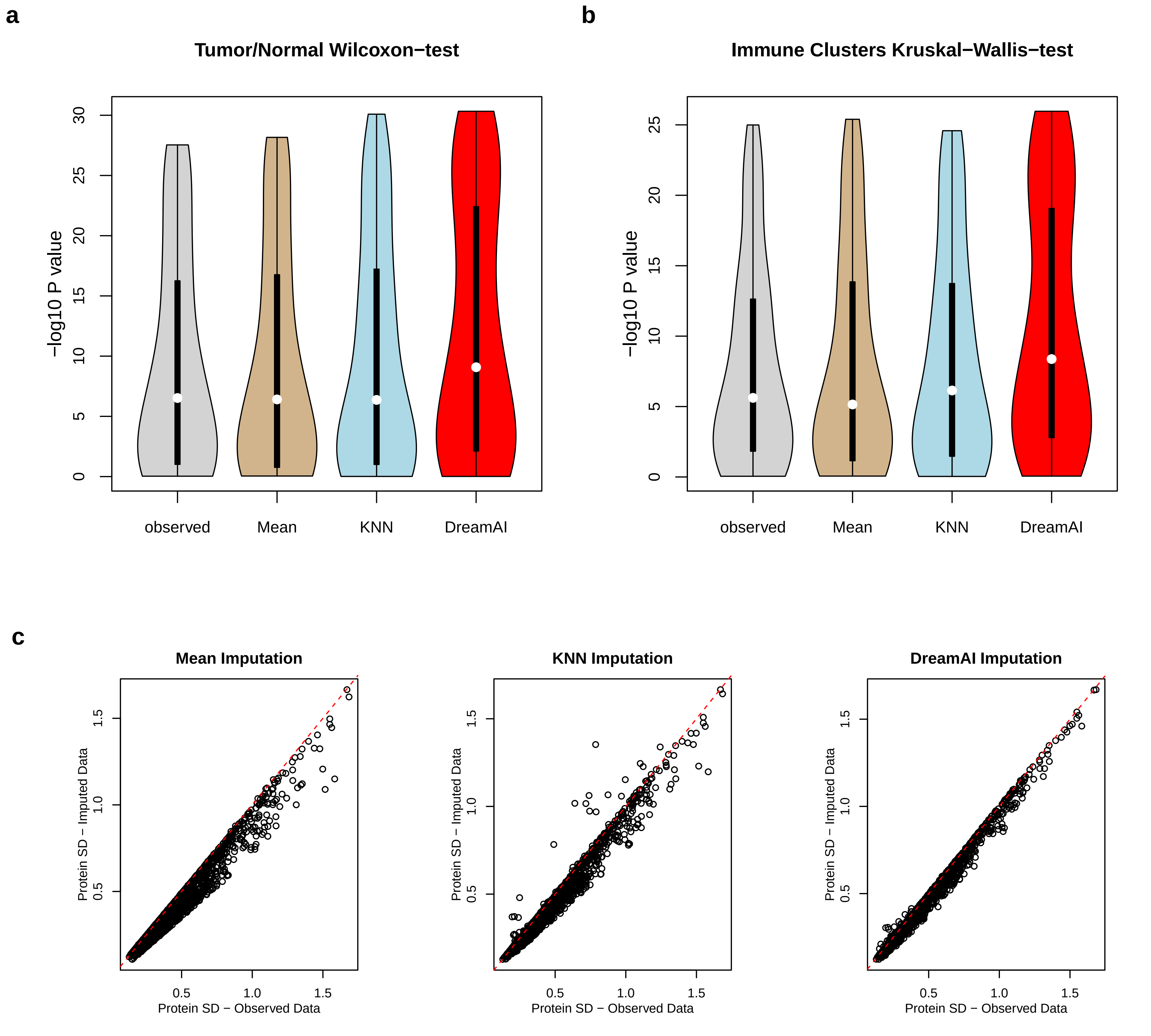
